# A unique small molecule pair controls the plant circadian clock

**DOI:** 10.1101/2020.05.25.113746

**Authors:** Takahiro N Uehara, Saori Takao, Hiromi Matsuo, Ami N. Saito, Eisuke Ota, Azusa Ono, Kenichiro Itami, Toshinori Kinoshita, Junichiro Yamaguchi, Norihito Nakamichi

## Abstract

Circadian clocks are the biological time keeping systems that coordinate genetic, metabolic, and physiological behaviors with the external day-night cycle. Previous studies have suggested possible molecular mechanisms for the circadian clock in *Arabidopsis thaliana* (Arabidopsis), but there might be additional mechanisms that have been hidden due to genetic redundancy.

A clock reporter line of Arabidopsis was screened against the 10,000 chemicals in the Maybridge Hitfinder10K chemical library, and a structure-activity relationship study of hit compounds was conducted. Clock mutants were treated with two of the small molecules to gain insight into their mode of action.

The screening identified 5-(3,4-dichlorophenyl)-1-phenyl-1,7-dihydro-4*H*-pyrazolo[3,4-*d*]pyrimidine-4,6(5*H*)-dione (**TU-892**) as a period lengthening molecule. From a structure-activity relationship study, we found that a molecule possessing 2,4-dichlorophenyl instead of a 3,4-dichlorophenyl group (**TU-923**) had period shortening activity. The period shortening activity of **TU-923** was reversed to a lengthening activity in double mutants lacking *PSEUDO-RESPONSE REGULATOR 9* (*PRR9*) and *PRR7* (*prr9-10 prr7-11*).

Our study provides a unique small molecule pair that regulates the pace of the clock in opposite ways, likely by targeting unknown factors. Small differences at the atomic level can reverse the period tuning activities. *PRR9* and *PRR7* are essential for the activity of **TU-923** in period shortening.

## Introduction

Circadian clocks are the biological timekeeping systems that govern the approximately 24-h rhythms of genetic, metabolic, physiological, and behavioral processes in most organisms. This oscillation allows organisms to predict and anticipate day-night changes in the environment. For instance, at dawn *Arabidopsis thaliana* (Arabidopsis) induces the expression of genes in the phenylpropanoid pathway that produce secondary metabolites capable of acting as phenolic sunscreens by absorbing ultraviolet and short-wavelength visible light (Harmer *et al.*, 2000). This phenomenon seems to be important for plants to anticipate and adapt to sunrise, which otherwise could result in photodamage to chloroplasts (Harmer *et al.*, 2000). Arabidopsis induces jasmonate defense hormones during the day for protection against damage from a herbivorous caterpillar, *Trichoplusia ni*, whose feeding activity is higher during the daytime and maximal at dusk (Goodspeed *et al.*, 2012). The plant clock contributes to fitness in 24 h day-night cycles even under laboratory conditions by coordinating the rhythms of many biological processes (Dodd *et al.*, 2005; Yerushalmi *et al.*, 2011). In addition to day-night anticipation, the clock is used to measure photoperiod for inducing flower meristem formation in many plants (Yanovsky & Kay, 2002). Consistently, genetic mutations in the clock are often found in some crop cultivars whose flowering times have changed due to human selection, thereby allowing the spread of these cultivars into areas or regions where the latitude or climate is different from the regions of origin for these crop plants (Nakamichi, 2015).

The molecular machinery at the core of the Arabidopsis circadian clock relies on transcription-translation feedback loops (TTFL) (Millar, 2016; Nohales & Kay, 2016; McClung, 2019). Evidence from genetic and biochemical studies have been taken to suggest the presence of Arabidopsis TTFLs. Among the TTFLs, there are some genetically redundant clock-associated genes (Millar, 2016; Nohales & Kay, 2016; McClung, 2019) due to whole-genome duplication followed by local duplication during evolution (Arabidopsis Genome Initiative, 2000). To investigate molecular mechanisms that potentially involve genetic redundancy, chemical genetic approaches have emerged. Chemical compounds used in these studies also overcome the lethality of gene mutations for studying gene function. Chemical compounds can also be applied in a dose-dependent, time-dependent, or growth stage-conditional manner, allowing stringent controls to be employed for each of the biological processes of interest (Uehara *et al.*, 2019).

Several studies have demonstrated that control of the animal clock can be achieved by small molecules, which will ultimately provide clock-modulator-based medicines (Hirota *et al.*, 2008; Isojima *et al.*, 2009; Hirota *et al.*, 2010; Hirota *et al.*, 2012; Tamai *et al.*, 2018; Oshima *et al.*, 2019). In plants, some natural or synthetic small molecules that modulate the clock have been reported. Endogenous sugars can entrain the Arabidopsis circadian clock (Haydon *et al.*, 2013). Prieurianin, also known as endosidin 1, was found to be a period shortening natural compound that perturbs actin cytoskeleton dynamics in Arabidopsis (Toth *et al.*, 2012). Molecules capable of destabilizing (latrunculin B and cytochalasin D) or stabilizing (jasplakinolide) actin also shorten the Arabidopsis circadian period (Toth *et al.*, 2012). Tetraethylammonium, a K+ channel blocker, shortens the circadian period in duckweed (*Lemna gibba* G3) (Kondo, 1990). Brevicompanine, a naturally occurring fungal compound, alters the expression of some clock-associated genes in Arabidopsis (de Montaigu *et al.*, 2017). Trichostatin A, a histone deacetylase inhibitor, causes a phase delay in clock-associated genes expressed during the evening in Arabidopsis (Perales & Mas, 2007).

Recently, we developed a high-throughput plant clock phenotyping system and found synthetic small molecules that lengthen the Arabidopsis circadian clock (Ono *et al.*, 2019; Uehara *et al.*, 2019). We have focused on molecules that change the circadian period since the robustness of period length stability in the midst of environmental fluctuations, such as temperature changes, is an important feature of circadian clocks. This property is striking from the point of view of the Arrhenius equation that describes the temperature dependence of chemical reactions (chemical reaction rates increase if temperatures increase). Generally, a 10°C increase causes a 2-fold increase in reaction rate (Q_10_ = 2). A chemical oscillation, the Belousov-Zhabotinsky reaction, follows the Arrhenius equation, and the period of oscillation is greatly shortened by increased temperatures (Bansagi *et al.*, 2009). The period of plant circadian clocks is not significantly altered by temperature changes, and the Q_10_ is far less than 2; this is called the temperature compensation of the period length (Salome *et al.*, 2010). Temperature compensation of circadian period length is found in the phosphorylation-dephosphorylation cycle of cyanobacterial KaiC clock proteins even *in vitro* (Nakajima *et al.*, 2005). If essential clock components are mutated, periods change duration. For example, both short- and long-period mutants were found for a *Drosophila* period gene (Bargiello *et al.*, 1984). Mutations in cyanobacterial *KaiC* resulted in short, long, or arrhythmic phenotypes, depending on the site of mutation (Ishiura *et al.*, 1998). Tuning the expression level of two clock-associated genes, *TIMING OF CAB EXPRESSION 1* (*TOC1*) or *ZEITLUPE* (*ZTL*), changes the period length, suggesting that period length is controlled by these genes (Mas *et al.*, 2003; Somers *et al.*, 2004). Thus, period change is a sign of perturbation or disorder of the clock.

PHA767491 was found to be a period lengthening molecule from the LOPAC library (Library of Pharmacologically Active Compounds that modulate a broad range of biological processes in mammals and microorganisms) that inhibited the casein kinase 1 family proteins in Arabidopsis (CKL proteins) (Uehara *et al.*, 2019). Through derivatization of PHA767491, we developed more a potent and selective Arabidopsis CKL inhibitor, AMI-331, that lengthens the period at higher nanomolar levels (Saito *et al.*, 2019). 3,4-Dibromo-7-azaindole (B-AZ) was also found as a period lengthening molecule from our in-house ‘ITbM’ chemical library that is enriched in plant hormone mimicking molecules, and B-AZ also inhibits CKLs (Ono *et al.*, 2019). These studies further found that treatment with these CKL inhibitors results in the accumulation of PSEUDO-RESPONSE REGULATOR 5 (PRR5) and TOC1 (known as PRR1) proteins, two transcriptional repressors in the clock genetic circuit. Consistently, CKL4 directly phosphorylates PRR5 and TOC1 for degradation (Uehara *et al.*, 2019). Finding a small molecule that modulates the circadian clock, followed by revealing the action mechanism of the hit molecule, makes it possible to reveal the molecular mechanisms underlying the clock. Therefore, chemical biology focusing on the plant clock provides considerable knowledge and chemical tools for developing and optimizing agrochemical use (de Montaigu *et al.*, 2017; Belbin *et al.*, 2019; Panter *et al.*, 2019; Uehara *et al.*, 2019).

To find additional small molecules capable of modulating the plant clock, we report here a high-throughput phenotypic screening using a different chemical library, the Maybridge Hitfinder 10K. 5-(3,4-Dichlorophenyl)-1-phenyl-*1H*-pyrazolo[3,4-*d*] pyrimidine-4,6(5*H,7H*)-dione (**TU-892**) was found to be a period lengthening molecule. A structure-activity relationship (SAR) study indicated that both the pyrazole ring and the pyrimidinedione moiety are essential. A molecule substituting a 2,4-dichlorophenyl moiety instead of a 3,4-dichlorophenyl group (**TU-923**), showed period-shortening activity. Our study provides evidence that a small difference at the atomic level can control the clock in opposite ways. Our results also suggest that there are crucial unknown factors involved in the clock mechanism that remain to be identified.

## Materials and methods

### Screening of small molecules that change the circadian period

Seeds of *Arabidopsis thaliana* accession Col-0 harboring *CCA1:LUC* (Nakamichi *et al.*, 2005) (a reporter construct that expresses luciferase with peak expression in the early morning) were sterilized and placed on half-strength Murashige-Skoog (MS) plates containing 0.25% (w/v) sucrose. Seeds were kept at 4°C in the dark for 2 days followed by transfer to 22°C with 12 h light (~70 µmol s^−1^ m^−2^) / 12 h dark conditions (LD). Four days after incubation, young seedlings were individually transferred with a dropper to a well of a 96-well plate. Seedlings were treated with small molecules from the Maybridge Hitfinder 10K library at a final concentration of 50 µM and 500 µM luciferin (120-05114, Wako). After an additional one-day incubation in LD conditions, the plates were read by a CL96 time-lapse luminescence detector (Churitsu, Toyoake, Japan). The circadian rhythm of the luminescent reporter was calculated as previously reported (Kamioka *et al.*, 2016). The molecule that changed the circadian period, [5-(3,4-dichlorophenyl)-1-phenyl-*1H*-pyrazolo[3,4-*d*]pyrimidine-4,6(5*H,7H*)-dione (**TU-892**)], was further confirmed by testing with different concentrations of the small molecule using *CCA1:LUC* and the evening reporter construct *TOC1:LUC* (Uehara *et al.*, 2019).

### *In vitro* phosphorylation assays of Arabidopsis CKLs

*In vitro* phosphorylation assays of Arabidopsis CKL4 and CKL1 were performed as previously reported (Uehara *et al.*, 2019). PHA767491 was purchased from Sigma Chemical Corp., dissolved in DMSO at a concentration of 10 mM, and stored at −20°C until use.

### Synthesis of TU-892 analogues

Synthesis of **TU-892** analogues is described in Supporting Information-Methods. **TU-892** analogues were dissolved in DMSO at a concentration of 10 mM and stored at room temperature. The effect of **TU-892** analogues on the circadian period length of Arabidopsis seedlings was tested as described above.

### TU-892 and TU-923 treatments of clock-period mutants

Clock-period mutants, *cca1-1 lhy-12 CCA1:LUC* (Kamioka *et al.*, 2016), *prr9-10 prr7-11 CCA1:LUC* (Nakamichi *et al.*, 2005), *prr7-11 prr5-11 CCA1:LUC* (Nakamichi *et al.*, 2005), and *prr5-11 toc1-2 CCA1:LUC* (Uehara *et al.*, 2019) were treated with **TU-892** and **TU-923** and the circadian rhythm assay was performed as described above.

## Results

### Screening synthetic small molecules that change the circadian period

To find small molecules capable of changing the circadian period of Arabidopsis seedlings, we conducted a high-throughput phenotypic screening using Maybridge Hitfinder 10K, a chemical library that is different from those used in our previous studies (Ono *et al.*, 2019; Uehara *et al.*, 2019). We monitored the circadian rhythm of the luminescent luciferase reporter driven by the *CIRCADIAN CLOCK-ASSOCIATED 1* (*CCA1*) promoter (*CCA1:luciferase* [*LUC*]), whose expression peaks in the early morning (Fig. **1a**). 5-(3,4-Dichlorophenyl)-1-phenyl-*1H*-pyrazolo[3,4-*d*]pyrimidine-4,6(5*H,7H*)-dione (**TU-892**) was found to be a period lengthening molecule (Fig. **1b**). **TU-892** lengthened the period of not only *CCA1:LUC* but also *TIMING OF CAB EXPRESSION 1:LUC* (*TOC1*:*LUC)*, whose luminescence peaks in the evening (Uehara *et al.*, 2019) in a dose-dependent fashion (Fig. **1c**,**d,e**). These results established **TU-892** as a period-lengthening molecule for the Arabidopsis circadian clock. Lower concentrations (25 µM) of TU-892 lengthen the circadian period about 2 h. Another parameter of the clock, amplitude, was not evaluated precisely in our test as previously reported (Ono *et al.*, 2019). Because we carefully selected seedlings of similar size, seedling size was unlikely the reason for the variable amplitude. We hypothesize that other factors, such as fluctuations in temperature during the assay, might have affected the amplitude. Variation in amplitude responding to environment fluctuations such as temperature is a general aspect of circadian rhythms (Murayama *et al.*, 2017). We also noticed that treatment with 100 µM **TU-892** caused bleaching of the seedlings, an indication of toxicity of this molecule at higher concentrations (Fig. S1). Since alterations of the circadian clock by genetic mutations do not cause lethality, **TU-892** at higher concentrations affects not only the circadian clock, but also essential physiological processes. Collectively, our result show that **TU-892** modulates an indispensable property of the clock, that is, the robustness of period length.

**Figure 1.**
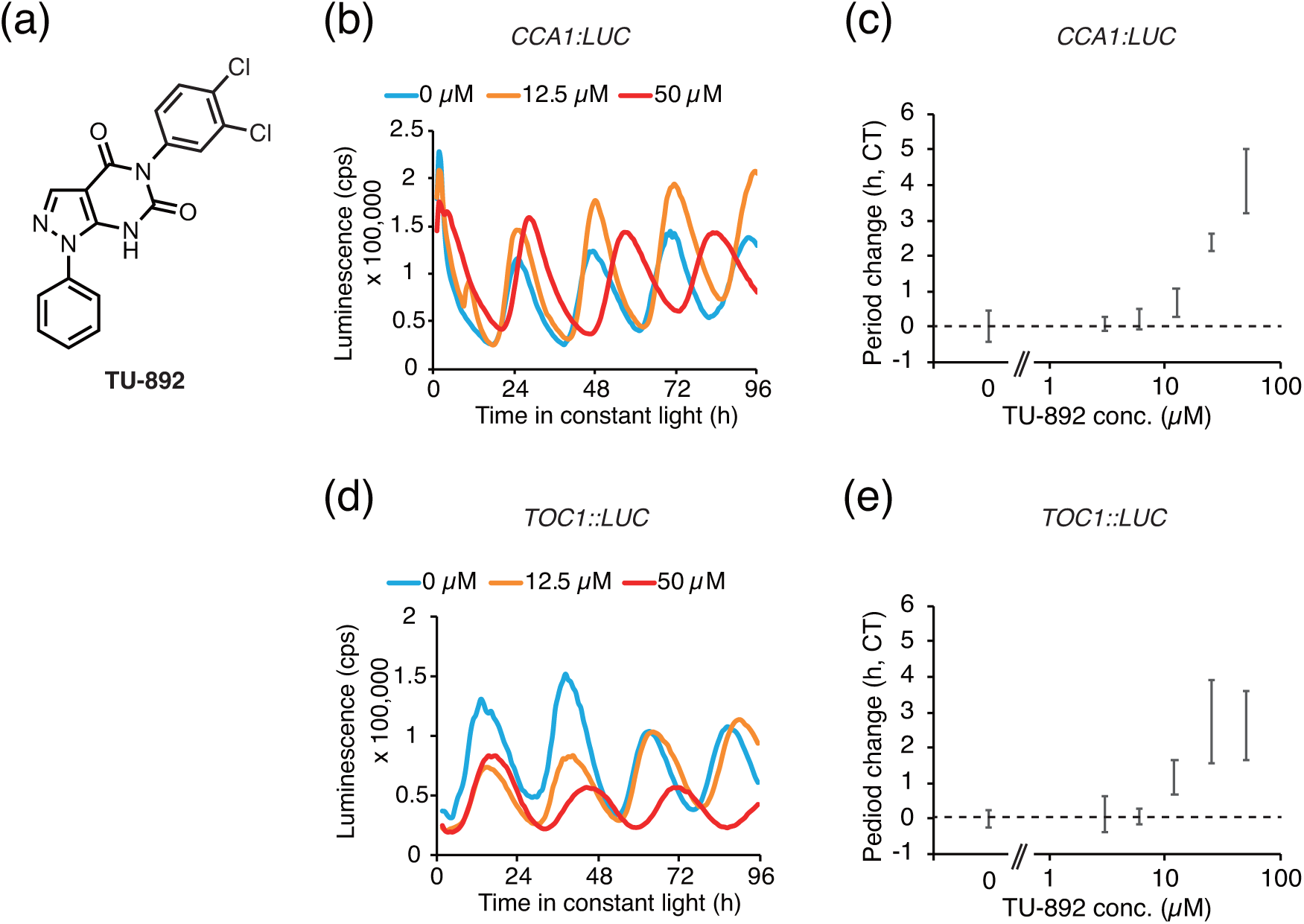
**TU-892** lengthens the Arabidopsis circadian period. (a) Chemical structure of **TU-892** (5-(3,4-dichlorophenyl)-1-phenyl-1,7-dihydro-4*H*-pyrazolo[3,4-*d*]pyrimidine-4,6(5*H*)-dione). Circadian luciferase reporter activity in Arabidopsis treated with **TU-892** *CCA1:LUC* (b), and *TOC1:LUC* (d). Increases in period length relative to the untreated (0 µM) control indicate a dose-response (c) and (e) (n = 7 or 8 for each concentration, with error bars indicating the standard deviation [S.D.]).

### TU-892 is not a CKL inhibitor

Given that three other period lengthening molecules (PHA767491, AMI-331, and B-AZ) inhibit CKL kinase activity (Ono *et al.*, 2019; Saito *et al.*, 2019; Uehara *et al.*, 2019), we examined whether **TU-892** also inhibits CKL. The CKL4 kinase activity for the model substrate casein was tested, because CKL4 kinase activity was the strongest among the purified CKL proteins (Uehara *et al.*, 2019). PHA767491 at concentrations of 4 to 100 µM strongly inhibited CKL4 kinase activity as previously reported (Uehara *et al.*, 2019), whereas the same concentration of **TU-892** did not inhibit CKL4 kinase activity at all (Fig. **2a**). PHA767491 also inhibited CKL1 kinase activity, whereas **TU-892** did not (Fig. **2b**), showing that **TU-892** is not a CKL inhibitor.

**Figure 2.**
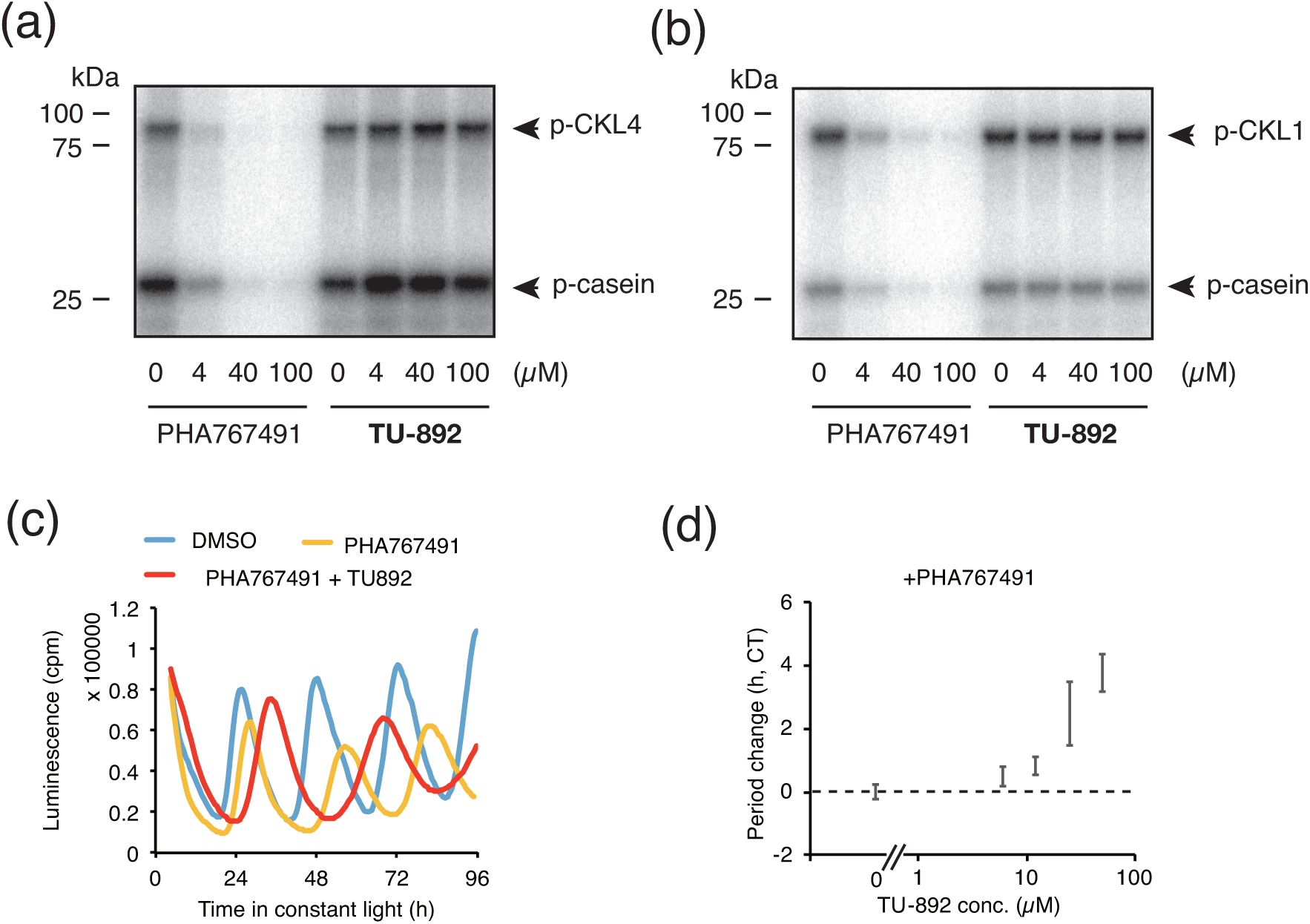
**TU-892** is not a CKL inhibitor. Autoradiography of *in vitro* CKL4 (a) and CKL1 (b) activities in the presence of **TU-892**. PHA767491 was used as a CKL inhibitor. (c) Luminescence from circadian luciferase reporter *CCA1:LUC* activity in Arabidopsis treated with 250 µM PHA767491 and 25 µM **TU-892**. (d) Increases in period length by **TU-892** relative to the untreated (0 µM) control in the presence of 250 µM PHA767491.

We also examined the interaction between CK1 inhibition and **TU-892** for period lengthening (Fig. 2**c,d**). PHA767491 treatment at a concentration of 250 µM lengthened the period by about 3 h, as previously reported (Uehara *et al.*, 2019). At a concentration of 250 µM, the effect of PHA767491 on the lengthening period is mostly saturated. If the mode of action of **TU-892** is dependent on CK1, **TU-892** should not lengthen the period in Arabidopsis treated with 250 µM PHA767491. Our results showed that **TU-892** lengthened the period in seedlings treated with 250 µM PHA767491 in a dose-dependent manner. This result suggested that **TU-892** lengthens the period independent of the inhibition of CK1 activity. The structure of **TU-892** is not similar to other molecules that potentially modulate clock parameters such as sugars, prieurianin, latrunculin B, cytochalasin D, jasplakinolide, tetraethylammonium, brevicompanine, and trichostatin A.

### Structure-activity relationship study of TU-892

To gain insight into the molecular mode of action of **TU-892** (**1a**) on period lengthening, we performed a structure-activity relationship (SAR) study. We initially synthesized **TU-892** and analogues as described in Supporting Information-Methods. We first tested whether the pyrimidinedione moiety is essential for period-lengthening activity (Fig. **3a**). Methyl substitution of pyrimidinedione had no period-lengthening activity (**2**). Replacement of the pyrimidinedione with a benzene ring had no activity, too (**3**). These results indicated that the pyrimidinedione is essential for period lengthening activity. Next, we evaluated the importance of the pyrazole ring. Two molecules replacing pyrazole with thiophene had quite weak or no period-lengthening activity (**4a,b**, Fig. **3b**). Note that these molecules lack the phenyl group on the pyrazole, suggesting that either the pyrazole or the phenyl group is essential for activity. Next, we substituted the phenyl group on the pyrazole. Substitution of the phenyl with methyl resulted in the loss of period-lengthening activity, suggesting the essentiality of the phenyl group (**1b**, Fig. **3c**). Replacement of the phenyl group with a *p*-tolyl (**1c**), *p*-nitrophenyl (**1e**), *p*-bromophenyl (**1f**), *m*-bromophenyl (**1g**), or *o*-bromophenyl (**1h**) group resulted in the loss of period-lengthening activity (Fig. **3c**). Replacement of the phenyl group with *p*-methoxyphenyl (**1d**) retained the activity. These results suggested that the phenyl group can be modified but there are limitations to the structural modifications that retain period-lengthening activity.

**Figure 3.**
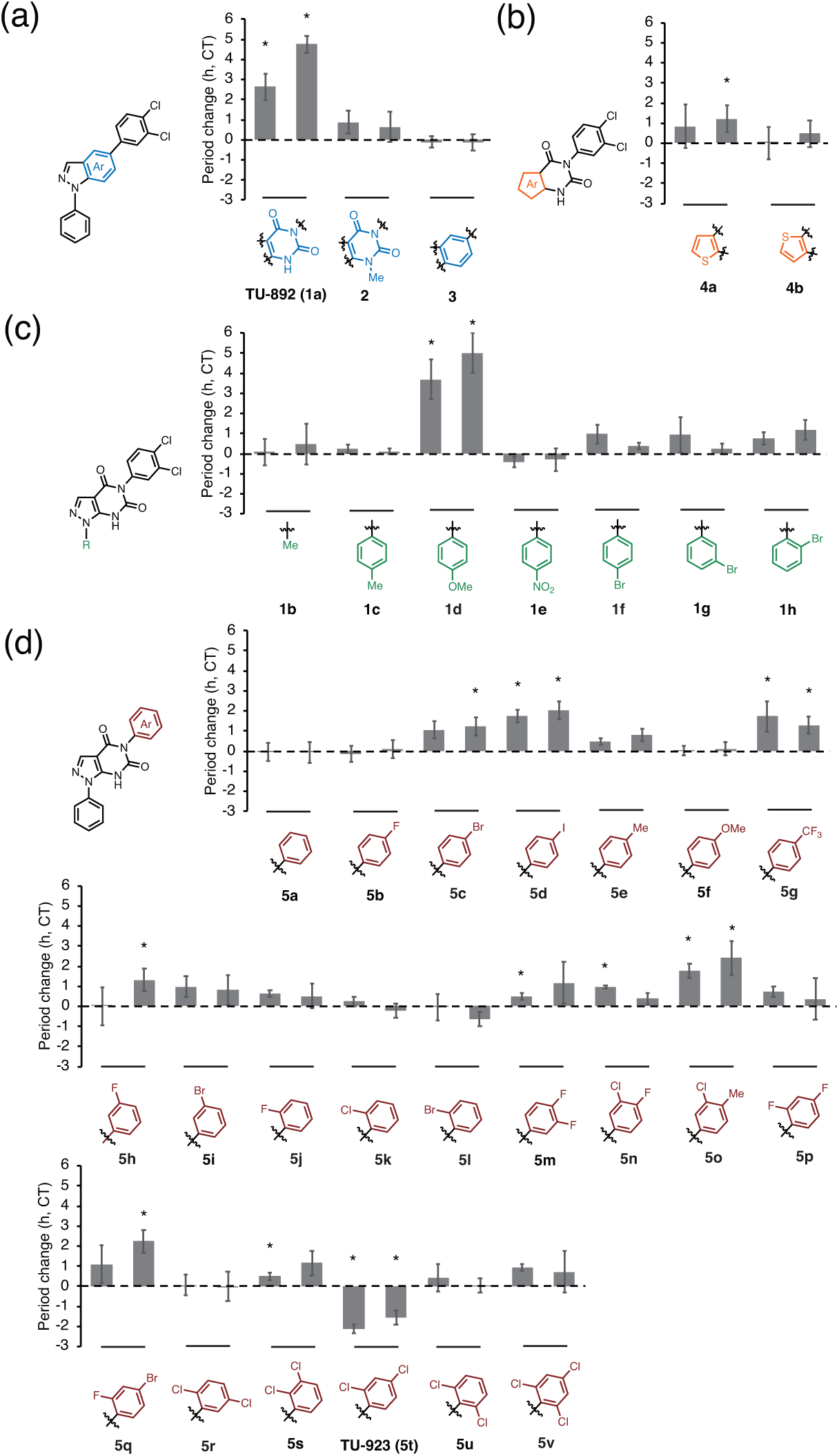
SAR study of **TU-892**. Circadian period changes after Arabidopsis seedling treatment with 25 (left) or 50 µM (right) **TU-892** analogues substituted at the pyrimidinedione (a), pyrazole (b), phenyl (c), or 3,4-dichlorophenyl (d) groups compared to solvent DMSO-treated samples are shown (n = 4, with S.D.). Asterisks indicate a significant change in circadian period compared to that of DMSO-treated samples (Student’s t-test *p* < 0.01).

We next examined the activity of derivatives modified at the 3,4-dichlorophenyl group on the pyrimidinedione (Fig. **3d**). Phenyl (**5a**), *p*-fluorophenyl (**5b**), *p*-tolyl (**5e**), *p*-methoxyphenyl (**5f**), *m*-bromophenyl (**5i**) *o*-fluorophenyl (**5j**), *o*-chlorophenyl (**5k**), *o*-bromophenyl (**5l**), 2,4-difluorophenyl (**5p**), 2,5-dichlorophenyl (**5r**), 2,6-dichlorophenyl (**5u**), and 2,4,6-trichlorophenyl (**5v**) derivatives had no period-lengthening activity. *p*-Bromophenyl (**5c**), *m*-fluorophenyl (**5h**), 3,4-difluorophenyl (**5m**), 3-chloro-4-fluorophenyl (**5n**), 2-fluoro-4-bromophenyl (**5q**), and 2,3-dichlorophenyl (**5s**) had low period-lengthening activities (significant changes were noted only with the 25 or 50 µM treatments). Reliable period-lengthening activities were found with the addition of *p*-iodophenyl (**5d**), *p*-trifluoromethyl (**5g**), and 3-chloro-4-methylphenyl (**5o**) (Fig. **3d**) analogues. Unexpectedly, period-shortening activity was found in the 2,4-dichlorophenyl (**5t**, **TU-923**) modified small molecule (Fig. **3d**).

Our SAR study indicated that both the pyrimidinedione and the pyrazole ring were essential for period-lengthening activity (Fig. **4a,b**). The phenyl group on the pyrazole was changeable. Interestingly, some modifications of the aryl group on the pyrimidinedione resulted in activity that shortened the circadian period. The period shortening activity of **TU-923** was further examined (Fig. **4c,d**). We found that **TU-923** shortened the period of both the morning (*CCA1:LUC*) and evening (*TOC1:LUC*) clock reporters, confirming that **TU-923** shortens the clock period. **TU-923** was also cytotoxic at a higher concentration (200 µM, Fig. S2).

**Figure 4.**
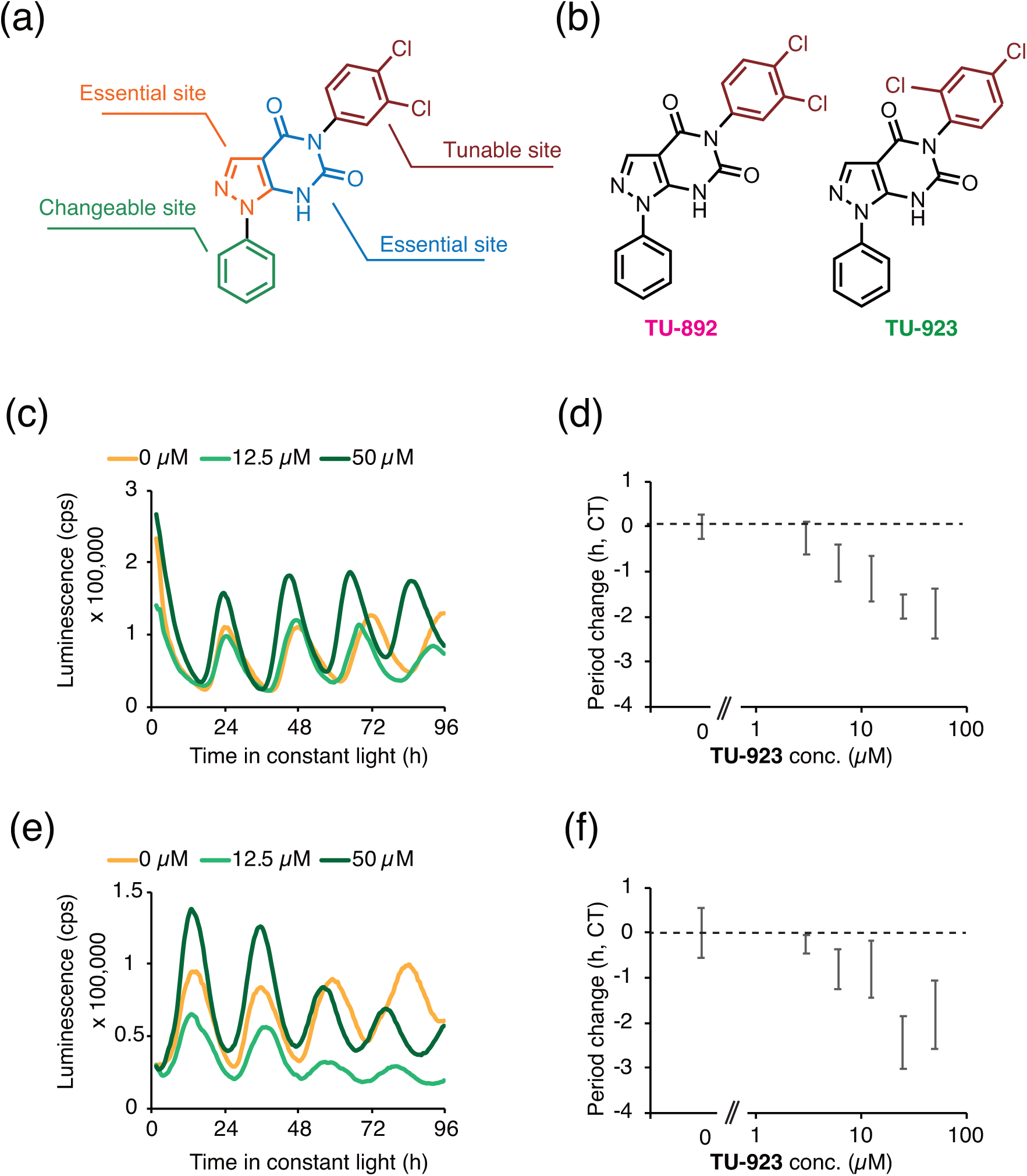
**TU-923** shortens the circadian period. (a) Summary of results from a SAR study of **TU-892**. (b) Structures of **TU-892** and **TU-923** (5-(2,4-dichlorophenyl)-1-phenyl-1,7-dihydro-4*H*-pyrazolo[3,4-*d*]pyrimidine-4,6(5*H*)-dione). Luminescence of circadian luciferase reporters *CCA1:LUC* (c) or *TOC1:LUC* (d) in Arabidopsis treated with **TU-923**. Period length changes relative to the untreated (0 µM) controls of *CCA1:LUC* (e) or *TOC1:LUC* (f) (n = 5 to 8 for each concentration, with error bars for S.D.).

### TU-923 lengthens the circadian period in *prr9 prr7* mutants

The effects of **TU-892** and **TU-923** on some clock-period mutants were also examined to help determine the action mechanisms of these molecules. We hypothesized that mutants impaired in a crucial gene associated with the mode of action of these molecules may experience a decrease in period modulating activities when treated with these small molecules. The short period mutants (*cca1-1 lhy-11* double mutants, *prr5-11 toc1-2* double mutants, or *prr7-11 prr5-11* double mutants) and a long period mutant (*prr9-10 prr7-11* double mutants) were treated with **TU-892** or **TU-923** (Fig. **5a**). In the wild type, 25 µM **TU-892** and **TU-923** lengthened and shortened the circadian period, respectively, as described above. **TU-892** lengthened the period in all of the clock mutants, indicating that **TU-892** does not require these genes for period lengthening. In contrast, the effect of **TU-923** diverged among the clock mutants. **TU-923** shortened the period in the *cca1-1 lhy-12* and *prr7-11 prr5-11* mutants, did not alter the period in *prr5-11 toc1-2*, and lengthened the period in *prr9-10 prr7-11*. To validate the period-lengthening activity of **TU-923** in *prr9-10 prr7-11*, we analyzed the mutants during continuous treatment with different concentrations of **TU-923** (Fig. **5b**). The results indicated that **TU-923** lengthened the period in these mutants in a dose-dependent fashion. The period lengthening effect of **TU-923** in *prr9-10 prr7-11* was about 4 - 8 h [Circadian Time (CT)-corrected] at concentrations of 25 to 50 µM. These results suggested that the period shortening activity of **TU-923** is required for the functions of *PRR9* and *PRR7*.

**Figure 5.**
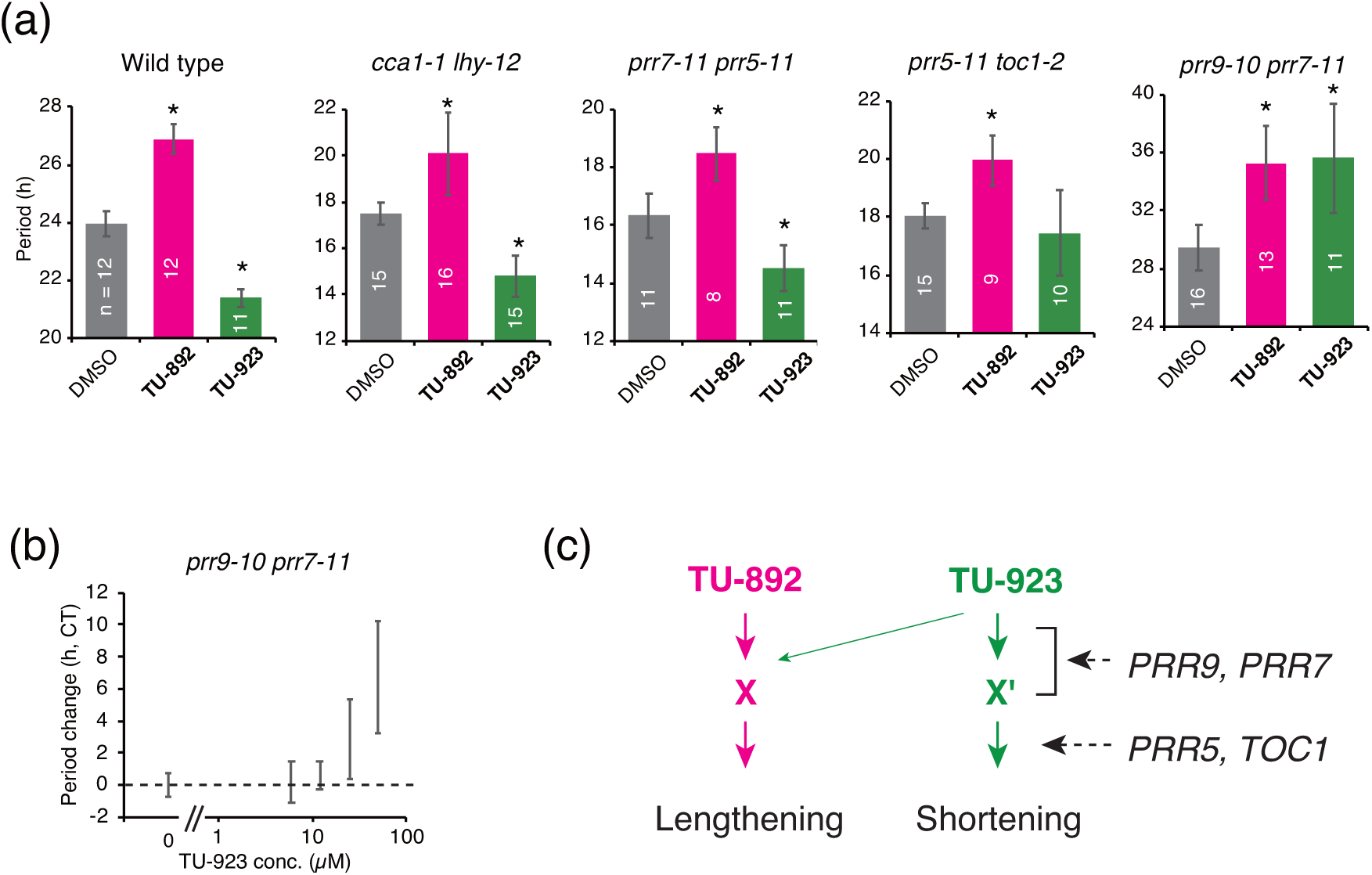
Effect of **TU-892** and **TU-923** on clock period mutants. (a) Effect of 25 µM **TU-892** and **TU-923** on period length of the *cca1-1 lhy-12*, *prr7-11 prr5-11*, *prr5-11 toc1-2*, and *prr9-10 prr7-11* mutants. Asterisks indicate significant differences in period length compared to that of the control (DMSO treatment) (Tukey-HSD test *p* < 0.05). Error bars indicate the S.D. (b) Period length change due to **TU-923** treatment relative to the untreated (0 µM) control of *prr9-10 prr7-11* (n = 5-11, with error bars for S.D.). All experiments were performed twice with similar results. (c) Proposed action mechanism for **TU-892** and **TU-923**. **TU-892** and **TU-923** target X and X’ (an X homologue), respectively. *PRR9* and *PRR7* participate in the interaction between **TU-923** and X’ or induce or activate X’. **TU-923** may change the target to X, if *PRR9* and *PRR7* are mutated. *PRR5* and *TOC1* are possibly implicated downstream of X’.

## Discussion

By a combined approach using a large-scale phenotypic screening and a SAR study, we found a period lengthening molecule, **TU-892**, and a period shortening molecule, **TU-923**, both of which have a similar chemical structure. Our study clearly indicated that **TU-892** is not a CKL inhibitor, suggesting the presence of a new pharmacologically tunable point for clock regulation. Importantly, reversing the direction of period length has not been achieved by any of the small molecule analogues of kinase inhibitors such as longdaysin and PHA767491 in animals and plants (Hirota *et al.*, 2010; Lee *et al.*, 2019; Saito *et al.*, 2019; Uehara *et al.*, 2019). Longdaysin and PHA767491 were proposed to inhibit CK1 kinase by binding to the ATP-binding pocket. Generally, it is quite difficult to make kinase activators that bind to the ATP-binding pocket, suggesting that TU-892 and TU-923 are unlikely to be competitive inhibitors of ATP.

A SAR study of the period lengthening molecule KL001, a modulator of the mammalian clock component cryptochrome, revealed a period shortening molecule (Oshima *et al.*, 2015). We speculated that **TU-892** and its analogue **TU-923** lengthens and shortens the circadian period, respectively, by controlling critical components of the Arabidopsis circadian clock. **TU-892** and **TU-923** structures differ only in the position of chloride in the dichlorophenyl moeity. 3,4-Dichlorophenyl can spin around on the bond between the dichlorophenyl group and the pyrimidine in **TU-892**, whereas 2,4-dichlorophenyl in **TU-923** cannot spin due to steric hindrance of the 2-chloro modification of the pyrimidine phenyl group. This structural difference may cause a difference in affinity or preference for binding to target proteins. Insight into the structural differences between **TU-892** and **TU-923** as well as the activity of **TU-923** in *prr9-10 prr7-11* will help reveal the full action mechanisms of these molecules for clock control.

Although the actual targets of **TU-892** and **TU-923** were not revealed in this study, we propose a model for the action mechanisms of **TU-892** and **TU-923** in period tuning (Fig. **5c**). Due to their structural similarity, **TU-892** and **TU-923** likely target very similar proteins or other endogenous factors (X and X’ in Fig. **5c**). The functions of X and X’ for period tuning are opposite. *PRR9* and *PRR7* participate in association with **TU-923** and X’ or induce or activate X’. T**U-923** becomes bound to X if *PRR9* and *PRR7* are mutated. *PRR5* and *TOC1* are implicated downstream of **TU-923** function, because the period of **TU-923** treatment in the *prr5-11 toc1-2* mutant was similar to the control experiment.

Since the plant clock regulates diverse biological processes including photoperiodic flowering time regulation and the drought stress response, both regarded as crucial traits for plant breeding programs, it should be possible to use clock modulators as agrochemicals. Unfortunately, however, at high concentrations **TU-892** and **TU-923** also have harmful effects on seedling growth as well as period-changing activities (Figs. S1, S2). Further derivatizations of **TU-892** and **TU-923** may provide molecules that will be useful as agrochemicals.

## Acknowledgements

We thank Drs H. Kasahara (Tokyo University for Agriculture and Technology), M. Kubo (Kumamoto University), and M. Hasebe (National Institute for Basic Biology, Okazaki, Aichi, Japan) for providing the chemical library Hitfinder 10K, and Dr. A. Sato and Ms. N. Kato for maintaining the TU-892 analogues. We thank Drs T. Kondo, K. Miwa, Y. Kitayama, T. Matsuo, H. Shinohara, and Y. Matsubayashi (Nagoya University) for helpful discussions. This work is supported in part by, a Grant-in-Aid for Scientific Research (KAKENHI) from the Japan Society for the Promotion of Science (18H02136) to NN, and a Grant-in-Aid for Scientific Research on Innovative Areas 18H04428 to J.Y., and 15H05956 to T.K. and 20H05411 to N.N. ITbM is supported by the World Premier International Research Center (WPI) Initiative, Japan.

## Author Contributions

TNU, ANS, EO, KI, and JY synthesized small molecules. ST conducted the chemical screening. HM and NN analyzed circadian assays. AO analyzed in vitro CK1 assays. KI and TK supervised the project. JY and NN conceptualized and wrote the paper.

## Supporting Information

Method: Synthesis Strategy for TU-892 and Analogues

Fig. S1 Treatment with a high concentration of TU-892 resulted in leaf bleaching.

Fig. S2 Treatment with a high concentration of TU-923 resulted in leaf bleaching.

**Fig. S1.**
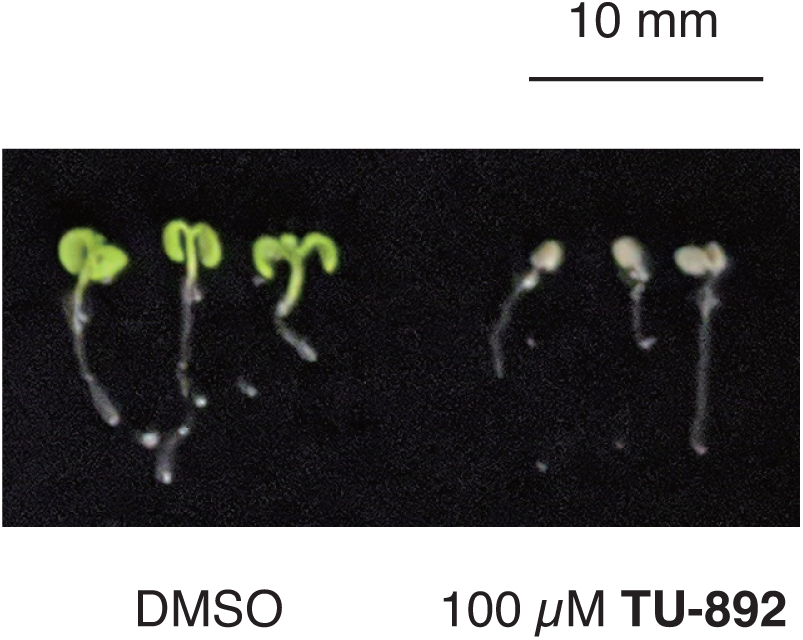
Treatment with a high concentration of **TU-892** resulted in leaf bleaching. Individual seedlings in a well of a 96-well plate were treated with 100 µM **TU-892** for 1 week. Cotyledons were bleached by the treatment.

**Fig. S2.**
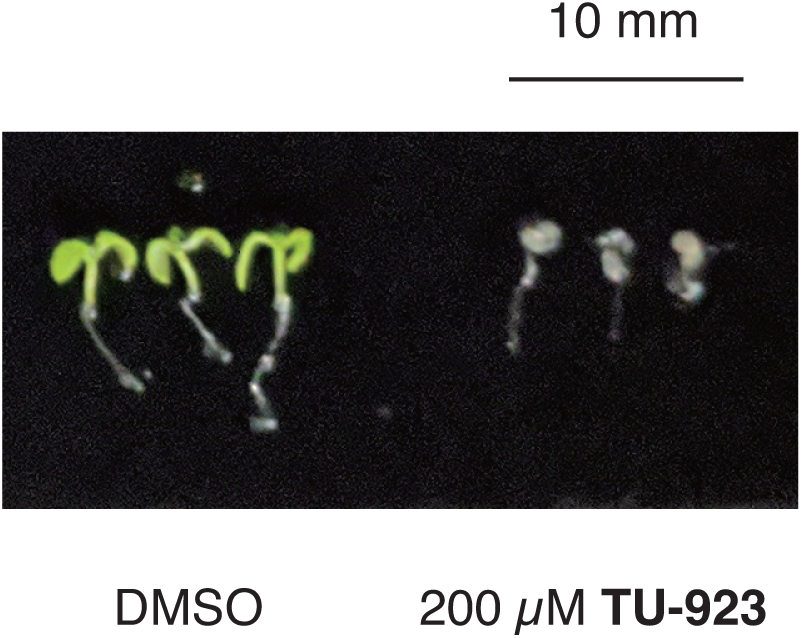
Treatment with a high concentration of **TU-923** resulted in leaf bleaching. Individual seedlings in a well of a 96-well plate were treated with 200 µM **TU-923** for 1 week. Cotyledons were bleached by the treatment.

