## supplemental text, method for "A unique small molecule pair controls the plant circadian clock"

**Synthesis of TU-892 Analogues**

**1. General**

Unless otherwise noted, all reactants or reagents including dry solvents were obtained from commercial suppliers and used as received. Ethyl 2-aminothiophene-3-carboxylate (**S1i**)[1] was synthesized according to procedures reported in the literature. Unless otherwise noted, all reactions were performed with dry solvents under an atmosphere of nitrogen in flame-dried glassware, using standard vacuum-line techniques. All work-up and purification procedures were carried out with reagent-grade solvents in air.

Analytical thin-layer chromatography (TLC) was performed using E. Merck silica gel 60 $F_{254}$ precoated plates (0.25 mm). The developed chromatogram was analyzed by UV lamp (254 nm). Flash column chromatography was performed with E. Merck silica gel 60 (230–400 mesh). Preparative thin-layer chromatography (PTLC) was performed using Wako-gel® B5-F silica coated plates (0.75 mm) prepared in our laboratory. High-resolution mass spectra (HRMS) were obtained from a Thermo Fisher Scientific Exactive (ESI) instrument. Nuclear magnetic resonance (NMR) spectra were recorded on a JEOL JNM-ECA-600 ($^{1}$H 600 MHz, $^{13}$C 150 MHz) spectrometer or a JEOL ECA 600II spectrometer with Ultra COOL™ probe ($^{1}$H 600 MHz, $^{13}$C 150 MHz). Chemical shifts for $^{1}$H NMR are expressed in parts per million (ppm) relative to tetramethylsilane (δ 0.00 ppm) or DMSO-$d_6$ (δ 2.50 ppm). Chemical shifts for $^{13}$C NMR are expressed in ppm relative to $CDCl_3$ (δ 77.0 ppm) or DMSO-$d_6$ (δ 40.0 ppm). Data are reported as follows: chemical shift, multiplicity (s = singlet, d = doublet, dd = doublet of doublets, dt = doublet of triplets, t = triplet, td = triplet of doublets, quin = quintet, sept = septet, m = multiplet, brs = broad signal), coupling constant (Hz), and integration.

### 2. Synthetic Strategy of TU-892 and Analogues

**Synthesis of TU-892 Analogues 1 and 4**

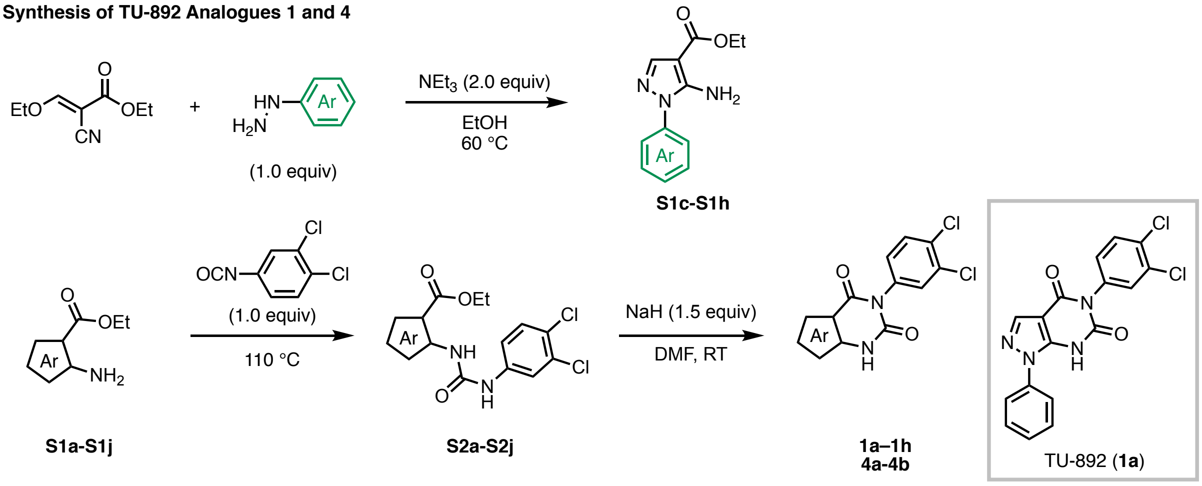


**Synthesis of TU-892 Analogue 2**

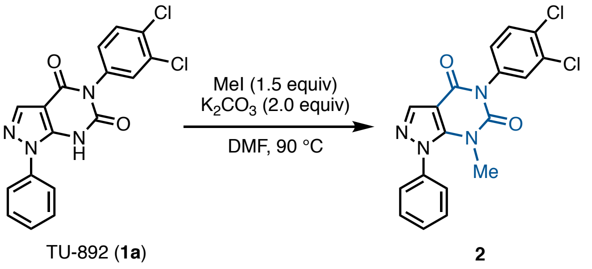


**Synthesis of TU-892 Analogue 3**

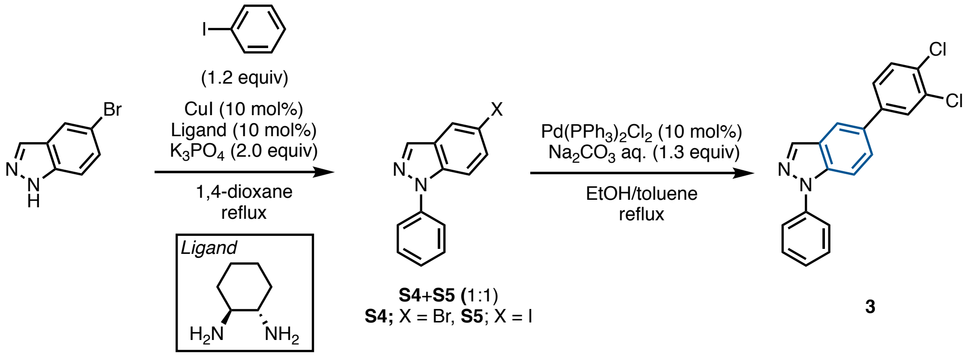


**Synthesis of TU-892 Analogues 5**

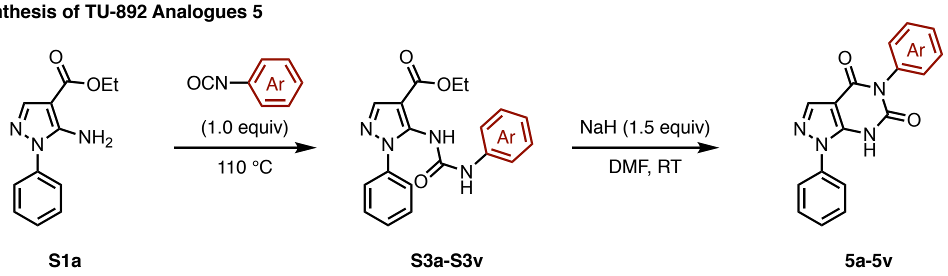

#### 3. Synthesis of TU-892 Analogues

##### 3.1 Synthesis of S1

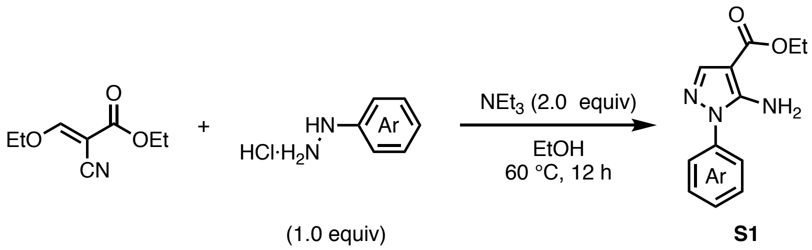


**S1**

##### General Procedure

To a solution of ethyl (*E*)-2-cyano-3-ethoxyacrylate (169.2 mg, 1.0 mmol) and arylhydrazine hydrochloride (1.0 mmol, 1.0 equiv) in EtOH (2.0 mL) was added triethylamine ($NEt_3$: 278.8 µL, 2.0 mmol, 2.0 equiv). After stirring at 60 °C for 12 h, the reaction was quenched with sat. $NH_4Cl$ aq. and extracted three times with EtOAc. The combined organic layer was washed with brine, dried over $Na_2SO_4$, and concentrated *in vacuo*. The residue was purified by flash column chromatography to afford **S1**.

**Ethyl 5-amino-1-(*p*-tolyl)-1*H*-pyrazole-4-carboxylate (S1c)**

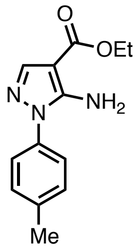


**S1c**

*p*-Tolylhydrazine hydrochloride was used as the arylhydrazine hydrochloride. Compound **S1c** was obtained as a yellow solid (201.1 mg, 82% yield). $^1H$ NMR (600 MHz, $CDCl_3$) δ 7.71 (s, 1H), 7.36 (d, $J$ = 8.4 Hz, 2H), 7.26 (d, $J$ = 8.4 Hz, 2H), 5.32 (brs, 2H), 4.27 (q, $J$ = 7.2 Hz, 2H), 2.38 (s, 3H), 1.34 (t, $J$ = 7.2 Hz, 3H); $^{13}C$ NMR (150 MHz, $CDCl_3$) δ 164.4, 148.9, 140.2, 138.0, 134.9, 130.1, 123.6, 95.8, 59.4, 20.9, 14.4; HRMS (ESI) $m/z$ = 268.1056 calcd for $C_{13}H_{15}N_3NaO_2$ $[M+Na]^+$, found: 268.1056.

**Ethyl 5-amino-1-(4-methoxyphenyl)-1*H*-pyrazole-4-carboxylate (S1d)**

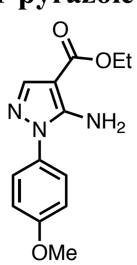


**S1d**

(4-Methoxyphenyl)hydrazine hydrochloride was used as the arylhydrazine hydrochloride. Compound **S1d** was obtained as a yellow solid (5.12 g, 98% yield). $^1H$ NMR (600 MHz,

CDCl3) δ 7.75 (s, 1H), 7.42 (d, $J$ = 9.0 Hz, 2H), 7.00 (d, $J$ = 9.0 Hz, 2H), 5.21 (brs, 2H), 4.30 (q, $J$ = 7.2 Hz, 2H), 3.85 (s, 3H), 1.36 (t, $J$ = 7.2 Hz, 3H); $^{13}$C NMR (150 MHz, $CDCl_3$) δ 164.6, 159.4, 149.1, 140.2, 130.3, 125.7, 114.8, 95.9, 59.6, 55.5, 14.5; HRMS (ESI) $m/z$ = 284.1006 calcd for $C_{13}H_{15}N_3NaO_3$ $[M+Na]^+$, found: 284.1006.

**Ethyl 5-amino-1-(4-nitrophenyl)-1*H*-pyrazole-4-carboxylate (S1e)**

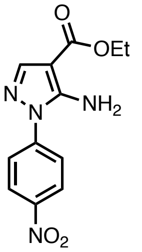


**S1e**

(4-Nitrophenyl)hydrazine hydrochloride was used as the arylhydrazine hydrochloride. Compound **S1e** was obtained as a yellow solid (7.11 g, 64% yield). $^{1}$H NMR (600 MHz, $CDCl_3$) δ 8.39 (d, $J$ = 9.0 Hz, 2H), 7.85–7.81 (m, 3H), 5.49 (brs, 2H), 4.32 (q, $J$ = 7.2 Hz, 2H), 1.38 (t, $J$ = 7.2 Hz, 3H); $^{13}$C NMR (150 MHz, $CDCl_3$) δ 164.3, 149.5, 146.2, 143.1, 141.9, 125.4, 123.0, 97.4, 60.0, 14.5; HRMS (ESI) $m/z$ = 275.0780 calcd for $C_{12}H_{11}N_4O_4$ $[M–H]^-$, found: 275.0780.

**Ethyl 5-amino-1-(4-bromophenyl)-1*H*-pyrazole-4-carboxylate (S1f)**

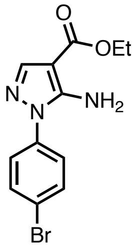


**S1f**

(4-Bromophenyl)hydrazine hydrochloride was used as the arylhydrazine hydrochloride. Compound **S1f** was obtained as a yellow solid (1.19 g, 77% yield). $^{1}$H NMR (600 MHz, $CDCl_3$) δ 7.78 (s, 1H), 7.64 (d, $J$ = 9.0 Hz, 2H), 7.44 (d, $J$ = 9.0 Hz, 2H), 5.30 (brs, 2H), 4.31 (q, $J$ = 7.2 Hz, 2H), 1.37 (t, $J$ = 7.2 Hz, 3H); $^{13}$C NMR (150 MHz, $CDCl_3$) δ 164.4, 149.0, 140.9, 136.6, 132.8, 125.1, 121.6, 96.4, 59.7, 14.4; HRMS (ESI) $m/z$ = 332.0005 calcd for $C_{12}H_{12}BrN_3NaO_2$ $[M+Na]^+$, found: 332.0005.

**Ethyl 5-amino-1-(3-bromophenyl)-1*H*-pyrazole-4-carboxylate (S1g)**

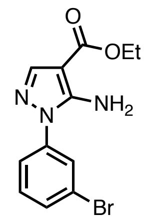


**S1g**

(3-Bromophenyl)hydrazine hydrochloride was used as the arylhydrazine hydrochloride. Compound **S1g** was obtained as a yellow solid (526.7 mg, 57% yield). $^1$H NMR (600 MHz, $CDCl_3$) δ 7.79 (s, 1H), 7.76 (t, *J* = 2.4 Hz, 1H), 7.55–7.50 (m. 2H), 7.38 (t, *J* = 8.4 Hz, 1H), 5.35 (brs, 2H), 4.31 (q, *J* = 7.2 Hz, 2H), 1.37 (t, *J* = 7.2 Hz, 3H); $^{13}$C NMR (150 MHz, $CDCl_3$) δ 164.3, 149.1, 140.9, 138.7, 130.9, 130.8, 126.6, 123.1, 121.8, 96.3, 59.7, 14.4; HRMS (ESI) *m/z* = 332.0005 calcd for $C_{12}H_{12}BrN_3NaO_2$ $[M+Na]^+$, found: 332.0004.

**Ethyl 5-amino-1-(2-bromophenyl)-1*H*-pyrazole-4-carboxylate (S1h)**

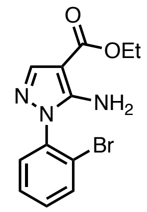


**S1h**

(2-Bromophenyl)hydrazine hydrochloride was used as the arylhydrazine hydrochloride. Compound **S1h** was obtained as a yellow solid (697.4 mg, 75% yield). $^1$H NMR (600 MHz, $CDCl_3$) δ 7.72 (s, 1H), 7.70 (dd, *J* = 7.8, 1.2 Hz, 1H), 7.43 (td, *J* = 7.8, 1.2 Hz, 1H), 7.39 (dd, *J* = 7.8, 1.8 Hz, 1H), 7.34 (td, *J* = 7.8, 1.8 Hz, 1H), 5.23 (brs, 2H), 4.25 (q, *J* = 7.2 Hz, 2H), 1.34 (t, *J* = 7.2 Hz, 3H); $^{13}$C NMR (150 MHz, $CDCl_3$) δ 164.1, 149.8, 140.5, 135.8, 133.6, 131.1, 129.8, 128.4, 121.9, 94.9, 59.3, 14.2; HRMS (ESI) *m/z* = 332.0005 calcd for $C_{12}H_{12}BrN_3NaO_2$ $[M+Na]^+$, found: 332.0003.

##### 3.2 Synthesis of 1 and 4

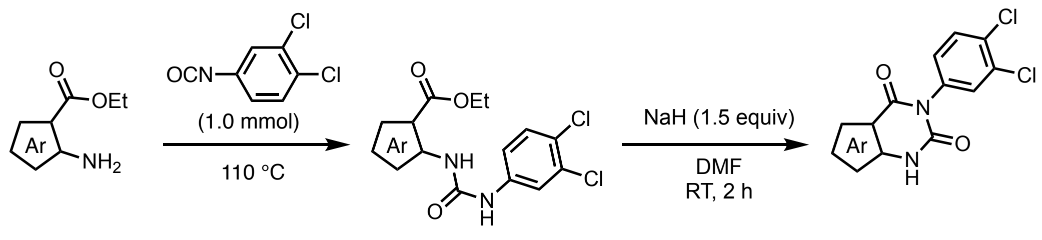


**S1**(1.0 equiv) **S2** **1** or **4**

##### General Procedure

A mixture of aryl carboxylate **S1** (1.0 equiv) and 1,2-dichloro-4-isocyanatobenzene (188.01 mg, 1.0 mmol) was stirred at 110 °C for several hours with monitoring reaction progress with TLC. After cooling to room temperature, the mixture was filtered, and the residue was washed with hexane/EtOAc (4:1). The residue was dissolved in DMF (1.0 mL) and sodium hydride (NaH: 60% dispersion in mineral oil, 60.0 mg, 1.5 mmol, 1.5 equiv) was added at 0 °C. After stirring at room temperature for 2 h, the reaction was quenched with sat. $NH_4Cl$ aq. and extracted three times with EtOAc. The combined organic layer was washed with brine, dried over $Na_2SO_4$, and concentrated *in vacuo*. The residue was purified by PTLC (hexane/EtOAc) to afford **1** or **4**.

**5-(3,4-Dichlorophenyl)-1-phenyl-1*H*-pyrazolo[3,4-*d*]pyrimidine- 4,6(5*H*,7*H*)-dione (1a: TU-892)**

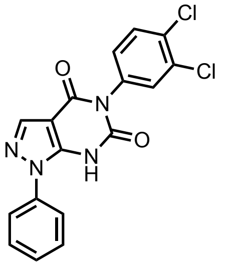

**TU-892**

Ethyl 5-amino-1-phenyl-1*H*-pyrazole-4-carboxylate (**S1a**) was used as the aryl carboxylate. TU-892 (**1a**) was obtained as a white solid (25.9 mg, 70% yield). $^1$H NMR (600 MHz, DMSO-$d_6$) δ 8.16 (s, 1H), 7.76 (d, $J$ = 9.0 Hz, 1H), 7.68 (d, $J$ = 2.4 Hz, 1H), 7.64 (d, $J$ = 7.8 Hz, 2H), 7.58 (t, $J$ = 7.8 Hz, 2H), 7.49 (t, $J$ = 7.8 Hz, 1H), 7.34 (dd, $J$ = 9.0, 2.4 Hz, 1H), one proton (NH) was not detected; $^{13}$C NMR (150 MHz, DMSO-$d_6$) δ 158.2, 151.6, 144.7, 138,1, 137.3, 136.8, 132.1, 131.5, 131.3, 131.2, 130.5, 129.9, 128.9, 124.7, 100.7; HRMS (ESI) *m/z* = 371.0108 calcd for $C_{17}H_9Cl_2N_4O_2$ [M–H]$^-$, found: 371.0103.

**5-(3,4-Dichlorophenyl)-1-methyl-1*H*-pyrazolo[3,4-*d*]pyrimidine-4,6(5*H*,7*H*)-dione (1b)**

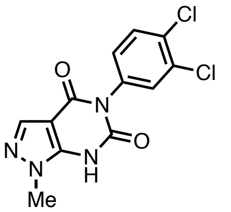

**1b**

Ethyl 5-amino-1-methyl-1*H*-pyrazole-4-carboxylate **S1b** was used as the aryl carboxylate. Compound **1b** was obtained as a white solid (16.8 mg, 26% yield). $^1$H NMR (600 MHz, DMSO-$d_6$) δ 7.90 (s, 1H), 7.74 (d, $J$ = 9.0 Hz, 1H), 7.68 (d, $J$ = 2.4 Hz, 1H), 7.33 (dd, $J$ = 9.0, 2.4 Hz, 1H), 3.82 (s, 3H), one proton (NH) was not detected; $^{13}$C NMR (150 MHz, DMSO-$d_6$) δ 157.9, 151.2, 144.0, 136.6, 136.5, 132.1, 131.5, 131.4, 131.1, 130.6, 99.6, 35.7; HRMS (ESI) *m/z* = 308.9946 calcd for $C_{12}H_7Cl_2N_4O_2$ [M–H]$^-$, found: 308.9948.

**5-(3,4-Dichlorophenyl)-1-(*p*-tolyl)-1*H*-pyrazolo[3,4-*d*]pyrimidine-4,6(5*H*,7*H*)-dione (1c)**

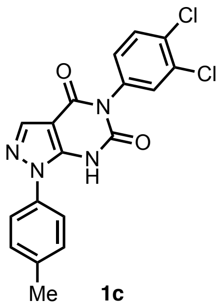

Me **1c**

Compound **S1c** was used as the aryl carboxylate. Compound **1c** was obtained as a white solid (17.5 mg, 18% yield). $^1$H NMR (600 MHz, DMSO-$d_6$) δ 12.50 (brs, 1H), 8.15 (s, 1H), 7.76 (d, *J* = 8.4 Hz, 1H), 7.69 (d, *J* = 2.4 Hz, 1H), 7.48 (d, *J* = 7.8 Hz, 2H), 7.38 (d, *J* = 7.8 Hz, 2H), 7.35 (dd, *J* = 8.4, 2.4 Hz, 1H), 2.40 (s, 3H), one proton (NH) was not detected; $^{13}$C NMR (150 MHz, DMSO-$d_6$) δ 158.1, 151.2, 143.9, 138.8, 137.9, 136.6, 134.7, 132.1, 131.5, 131.4, 131.2, 130.5, 130.4, 124.9, 100.6, 21.2; HRMS (ESI) *m/z* = 385.0259 calcd for $C_{18}H_{11}Cl_2N_4O_2$ [M–H]$^-$, found: 385.0262.

**5-(3,4-Dichlorophenyl)-1-(4-methoxyphenyl)-1,7-dihydro-4*H*-pyrazolo[3,4-*d*]pyrimidine-4,6(5*H*)-dione (1d)**

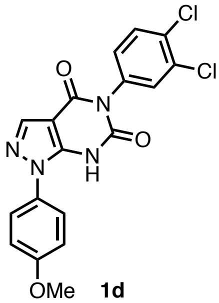

OMe **1d**

The reaction was performed at 60 °C. Compound S**1d** was used as the aryl carboxylate. Compound **1d** was obtained as a white solid (48.8 mg, 48% yield). $^1$H NMR (600 MHz, DMSO-$d_6$) δ 12.48 (brs, 1H), 8.12 (s, 1H), 7.75 (d, *J* = 8.4 Hz, 1H), 7.68 (d, *J* = 1.8 Hz, 1H), 7.51 (d, *J* = 9.0 Hz, 2H), 7.34 (dd, *J* = 8.4, 1.8 Hz, 1H), 7.11 (d, *J* = 9.0 Hz, 2H), 3.84 (s, 3H); $^{13}$C NMR (150 MHz, DMSO-$d_6$) δ 159.9, 158.1, 151.3, 144.2, 137.7, 136.7, 132.1, 131.5, 131.4, 131.2, 130.5, 130.0, 126.8, 115.0, 100.4, 56.0; HRMS (ESI) *m/z* = 401.0208 calcd for $C_{18}H_{11}Cl_2N_4O_3$ [M–H]$^-$, found: 401.0212.

**5-(3,4-Dichlorophenyl)-1-(4-nitrophenyl)-1*H*-pyrazolo[3,4-*d*]pyrimidine-4,6(5*H*,7*H*)-dion e (1e)**

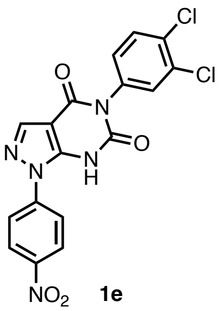

NO2 **1e**

Compound S**1e** was used as the aryl carboxylate. Compound **1e** was obtained as a white solid (18.5 mg, 26% yield). $^1$H NMR (600 MHz, DMSO-*d*$_6$) δ 8.46 (d, *J* = 9.0 Hz, 2H), 8.37 (d, *J* = 9.0 Hz, 2H), 8.04 (s, 1H), 7.68 (d, *J* = 8.4 Hz, 1H), 7.51 (d, *J* = 2.4 Hz, 1H), 7.21 (dd, *J* = 8.4, 2.4 Hz, 1H), one proton (NH) was not detected; $^{13}$C NMR (150 MHz, DMSO-*d*$_6$) δ 159.6, 156.0, 155.4, 144.9, 144.5, 139.0, 132.1, 131.1, 130.8, 130.6, 130.2, 125.3, 121.0, 100.3, one peak is missing due to overlapping; HRMS (ESI) *m/z* = 415.9953 calcd for $C_{17}H_8Cl_2N_5O_4$ $[M–H]^-$, found: 415.9953.

**1-(4-Bromophenyl)-5-(3,4-dichlorophenyl)-1*H*-pyrazolo[3,4-*d*]pyrimidine-4,6(5*H*,7*H*)-dione (1f)**

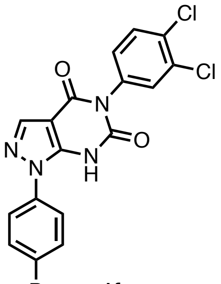

Br **1f**

Compound **S1f** was used as the aryl carboxylate. Compound **1f** was obtained as a white solid (69.8 mg, 62% yield). $^1$H NMR (600 MHz, DMSO-*d*$_6$) δ 8.18 (s, 1H), 7.83–7.73 (m, 3H), 7.67 (d, *J* = 1.8 Hz, 1H), 7.59 (d, *J* = 8.4 Hz, 2H), 7.33 (dd, *J* = 8.4, 1.8 Hz, 1H), one proton (NH) was not detected; $^{13}$C NMR (150 MHz, DMSO-*d*$_6$) δ 158.1, 151.4, 144.5, 138.4, 136.6, 136.4, 132.9, 132.0, 131.5, 131.4, 131.2, 130.5, 127.0, 122.0, 100.8; HRMS (ESI) *m/z* = 448.9208 calcd for $C_{17}H_8BrCl_2N_4O_2$ $[M–H]^-$, found: 448.9208.

**1-(3-Bromophenyl)-5-(3,4-dichlorophenyl)-1*H*-pyrazolo[3,4-*d*]pyrimidine-4,6(5*H*,7*H*)-dione (1g)**

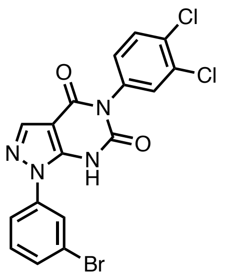

**1g**

Compound **S1g** was used as the aryl carboxylate. Compound **1g** was obtained as a white solid (51.2 mg, 39% yield). $^1$H NMR (600 MHz, DMSO-$d_6$) δ 8.15 (s, 1H), 7.95 (s, 1H), 7.78–7.71 (m, 2H), 7.68–7.61 (m, 2H), 7.52 (t, $J$ = 7.8 Hz, 1H), 7.31 (dd, $J$ = 7.8, 1.8 Hz, 1H), one proton (NH) was not detected; $^{13}$C NMR (150 MHz, DMSO-$d_6$) δ 157.9, 151.8, 145.9, 138.6, 138.0, 136.6, 131.7, 131.3, 131.2, 131.1, 130.9, 130.7, 130.1, 126.7, 123.1, 121.9, 100.4; HRMS (ESI) $m/z$ = 448.9208 calcd for $C_{17}H_8BrCl_2N_4O_2$ [M–H]$^-$, found: 448.9209.

**1-(2-Bromophenyl)-5-(3,4-dichlorophenyl)-1*H*-pyrazolo[3,4-*d*]pyrimidine-4,6(5*H*,7*H*)-dione (1h)**

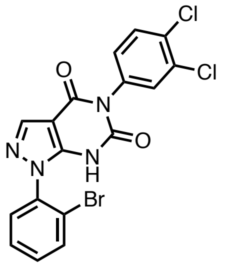

**1h**

Compound S**1h** was used as the aryl carboxylate. Compound **1h** was obtained as a white solid (59.2 mg, 45% yield). $^1$H NMR (600 MHz, DMSO-$d_6$) δ 8.20 (s, 1H), 7.88 (d, $J$ = 7.8 Hz, 1H), 7.76–7.73 (m, 2H), 7.63–7.52 (m, 3H), 7.38 (d, $J$ = 8.4 Hz, 1H), one proton (NH) was not detected; $^{13}$C NMR (150 MHz, DMSO-$d_6$) δ 158.0, 151.0, 145.2, 138.2, 136.4, 136.0, 133.9, 132.7, 132.1, 131.6, 131.5, 131.2, 130.9, 130.5, 129.4, 122.3, 99.8; HRMS (ESI) $m/z$ = 448.9208 calcd for $C_{17}H_8BrCl_2N_4O_2$ [M–H]$^-$, found: 448.9212.

**3-(3,4-Dichlorophenyl)thieno[2,3-*d*]pyrimidine-2,4(1*H*,3*H*)-dione (4a)**

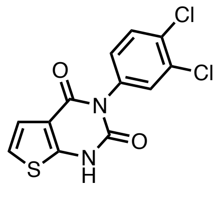

**4a**

Ethyl 2-aminothiophene-3-carboxylate (**S1i**) was used as the aryl carboxylate. Purification by PTLC ($CHCl_3$/MeOH = 20:1) afforded **4a** as a white solid (29.7 mg, 19% yield). $^1$H NMR

(600 MHz, DMSO-$d_6$) δ 12.41 (brs, 1H), 7.75 (d, $J$ = 8.4 Hz, 1H), 7.73 (d, $J$ = 2.4 Hz, 1H), 7.37 (dd, $J$ = 8.4, 2.4 Hz, 1H), 7.20 (d, $J$ = 5.4 Hz, 1H), 7.15 (d, $J$ = 5.4 Hz, 1H); $^{13}$C NMR (150 MHz, DMSO-$d_6$) δ 158.9, 152.0, 150.7, 136.4, 132.0, 131.53, 131.50, 131.2, 130.4, 122.8, 117.9, 115.5; HRMS (ESI) $m/z$ = 310.9449 calcd for $C_{12}H_5Cl_2N_2O_2S$ [M–H]$^-$, found: 310.9449.

**3-(3,4-Dichlorophenyl)thieno[3,2-*d*]pyrimidine-2,4(1*H*,3*H*)-dione (4b)**

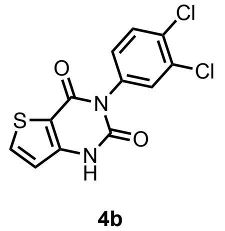

**4b**

Ethyl 3-aminothiophene-2-carboxylate (**S1j**) was used as the aryl carboxylate. Purification by PTLC ($CHCl_3$/MeOH = 20:1) afforded **4b** as a white solid (142.2 mg, 30% yield). $^1$H NMR (600 MHz, DMSO-$d_6$) δ 12.07 (brs, 1H), 8.13 (d, $J$ = 5.4 Hz, 1H), 7.77–7.73 (m, 2H), 7.39 (dd, $J$ = 8.4, 2.4 Hz, 1H), 6.98 (d, $J$ = 5.4 Hz, 1H); $^{13}$C NMR (150 MHz, DMSO-$d_6$) δ 158.6, 151.5, 145.9, 137.2, 136.3, 132.0, 131.5, 131.2, 130.5, 117.8, 111.7, one peak is missing due to overlapping; HRMS (ESI) $m/z$ = 310.9449 calcd for $C_{12}H_5Cl_2N_2O_2S$ [M–H]$^-$, found: 310.9452.

**3.3 Synthesis of 5-(3,4-Dichlorophenyl)-7-methyl-1-phenyl-1*H*-pyrazolo[3,4-*d*] -pyrimidine-4,6(5*H*,7*H*)-dione (2)**

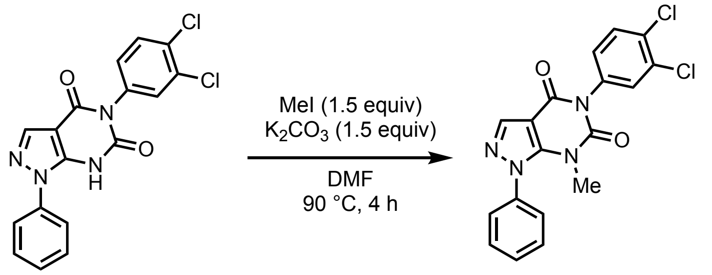


**TU-892 (1a)** **2**

To a solution of TU-892 (29.5 mg, 0.08 mmol) and $K_2CO_3$ (16.4 mg, 0.12 mmol, 1.5 equiv) in DMF (0.3 mL) was added iodomethane (7.4 µL, 0.12 mmol, 1.5 equiv). After stirring at 90 °C for 4 h, the reaction was quenched with sat. $NH_4Cl$ aq. and extracted three times with EtOAc. The combined organic layer was washed with brine, dried over $Na_2SO_4$, and concentrated *in vacuo*. The residue was purified by PTLC ($CHCl_3$/MeOH = 40:1) to afford **2** as a white solid (21.6 mg, 70% yield). $^1$H NMR (600 MHz, $CDCl_3$) δ 8.13 (s, 1H), 7.60–7.56 (m, 4H), 7.53–7.50 (m, 2H), 7.39 (d, $J$ = 2.4 Hz, 1H), 7.13 (dd, $J$ = 7.8, 2.4 Hz, 1H), 3.15 (s, 3H); $^{13}$C NMR (150 MHz, DMSO-$d_6$) δ 156.8, 151.3, 144.9, 138.7, 137.6, 136.4, 131.5, 131.3, 130.9, 130.3, 129.9, 129.5, 127.8, 101.4, 32.7, one peak is missing due to overlapping; HRMS (ESI) $m/z$ = 409.0230 calcd for $C_{18}H_{12}Cl_2N_4NaO_2$ [M+Na]$^+$, found: 409.0229.

##### 3.4 Synthesis of 5-(3,4-Dichlorophenyl)-1-phenyl-1*H*-indazole (3)

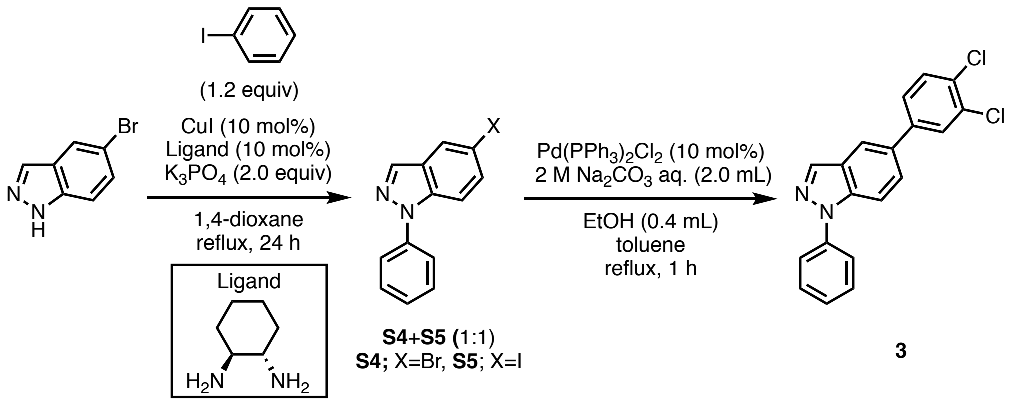


**S4**+**S5** (1:1)
**S4**; X=Br, **S5**; X=I

**3**

To a 200-mL three-necked flask, containing a magnetic stirring bar, were added CuI (57.0 mg, 0.3 mmol, 10 mol%), (1*S*,2*S*)-cyclohexane-1,2-diamine (0.41 mL, 0.3 mmol, 10 mol%), iodobenzene (0.40 mL, 3.6 mmol, 1.2 equiv), 5-bromo-1*H*-indazole (591.1 mg, 3.0 mmol, 1.0 equiv), and 1.4-dioxane (30 mL). The the mixture was refluxed for 24 h. After cooling to room temperature, the mixture was passed through a short silica gel pad with EtOAc as an eluent and then concentrated *in vacuo*. The residue was purified by flash column chromatography (hexane/EtOAc = 20:1) to afford 5-bromo-1-phenyl-1*H*-indazole (**S4**) and 5-iodo-1-phenyl-1*H*-indazole (**S5**) as a mixture. To a 30-mL two-necked flask, containing a magnetic stirring bar, were added $Pd(PPh_3)_2Cl_2$ (35.0 mg, 0.05 mmol, 10 mol%), 2 M $Na_2CO_3$ aq. (2.0 mL), EtOH (0.4 mL), the mixture (**S4**+**S5**: 148.3 mg, 0.5 mmol, 1.0 equiv), and toluene (2.0 mL). The mixture was refluxed for 1 h. After cooling to room temperature, the mixture was passed through a short silica gel pad with EtOAc as an eluent and then dried over $Na_2SO_4$. The mixture was filtered and concentrated *in vacuo*. The residue was purified by PTLC (hexane/EtOAc = 10:1, then $CHCl_3$) to afford **3** as a yellow solid (86.8 mg, 51% yield). $^1$H NMR (600 MHz, $CDCl_3$) δ 8.25 (s, 1H), 7.93 (m, 1H), 7.80 (d, $J$ = 8.4 Hz, 1H), 7.74 (d, $J$ = 7.8 Hz, 2H), 7.72 (d, $J$ = 1.8 Hz, 1H), 7.60 (dd, $J$ = 8.4, 1.8 Hz, 1H), 7.56 (t, $J$ = 7.8 Hz, 2H), 7.51 (d, $J$ = 8.4 Hz, 1H), 7.46 (dd, $J$ = 8.4, 1.8 Hz, 1H), 7.39 (t, $J$ = 7.8 Hz, 1H); $^{13}$C NMR (150 MHz, $CDCl_3$) δ 141.2, 139.9, 138.4, 135.8, 132.9, 132.6, 131.2, 130.7, 129.5, 129.1, 126.9, 126.6, 126.5, 125.9, 122.7, 119.5, 111.0; HRMS (ESI) $m/z$ = 339.0450 calcd for $C_{19}H_{13}Cl_2N_2$ $[M+H]^+$, found: 339.0450.

##### 3.5 Synthesis of Compound 5

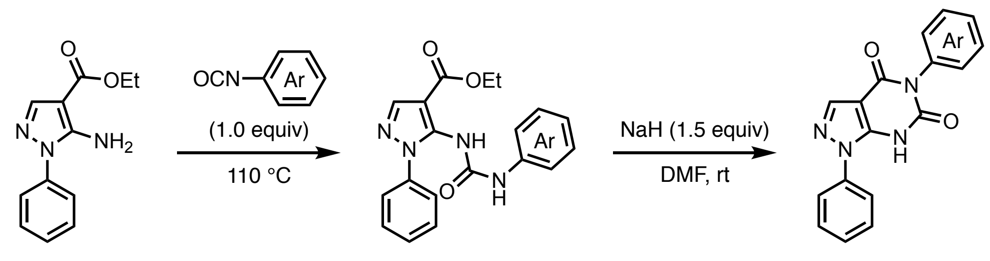


**S1a** **S3** **5**

##### General Procedure

A mixture or ethyl 5-amino-1-phenyl-1*H*-pyrazole-4-carboxylate (**S1a**: 233 mg, 1.0 mmol) and aryl isocyanate (1.0 equiv) was stirred at 110 °C for several hours with monitoring reaction progress with TLC. After cooling to room temperature, the mixture was filtered and the residue was washed with hexane/EtOAc (4:1). The residue was dissolved in DMF (1.0 mL) and sodium hydride (NaH: 60% dispersion in mineral oil, 60.0 mg, 1.5 mmol, 1.5 equiv) was added at 0 °C. After stirring at room temperature for 2 h, the reaction was quenched with sat. $NH_4Cl$ aq. and extracted three times with EtOAc. The combined organic layer was washed with brine, dried over $Na_2SO_4$, and concentrated *in vacuo*. The residue was purified by PTLC to afford **5**.

##### 1,5-Diphenyl-1*H*-pyrazolo[3,4-*d*]pyrimidine-4,6(5*H*,7*H*)-dione (5a)

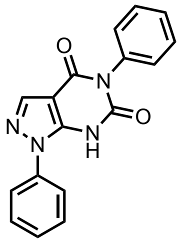

**5a**

Isocyanatobenzene was used as the aryl isocyanate. Compound **5a** was obtained as a white solid (38.0 mg, 50% yield). $^1$H NMR (600 MHz, DMSO-$d_6$) δ 12.44 (brs, 1H), 8.15 (s, 1H), 7.64 (d, $J$ = 7.8 Hz, 2H), 7.60–7.55 (m, 2H), 7.53–7.45 (m, 3H), 7.41 (t, $J$ = 7.8 Hz, 1H), 7.27 (d, $J$ = 7.8 Hz, 2H); $^{13}$C NMR (150 MHz, DMSO-$d_6$) δ 158.4, 151.6, 144.1, 138.1, 137.2, 136.5, 129.9, 129.8, 129.3, 129.0, 128.4, 124.9, 100.8; HRMS (ESI) $m/z$ = 303.0882 calcd for $C_{17}H_{11}N_4O_2$ [M–H]$^-$, found: 303.0882.

##### 5-(4-Fluorophenyl)-1-phenyl-1*H*-pyrazolo[3,4-*d*]pyrimidine-4,6(5*H*,7*H*)-dione (5b)

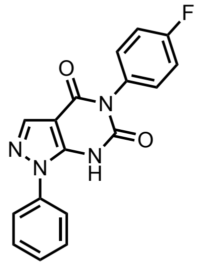

**5b**

1-Fluoro-4-isocyanatobenzene was used as the aryl isocyanate. Compound **5b** was obtained as a white solid (19.7 mg, 61% yield). $^{1}$H NMR (600 MHz, DMSO-*d*$_6$) δ 12.47 (brs, 1H), 8.12 (s, 1H), 7.68 (d, *J* = 7.8 Hz, 2H), 7.56 (t, *J* = 7.8 Hz, 2H), 7.47 (t, *J* = 7.8 Hz, 1H), 7.34–7.26 (m, 4H); $^{13}$C NMR (150 MHz, DMSO-*d*$_6$) δ 161.9 (d, $J_{C\text{–}F}$ = 241.2 Hz), 158.6, 152.1, 145.2, 138.0, 137.5, 133.0, 131.8 (d, $J_{C\text{–}F}$ = 8.6 Hz), 129.9, 128.7, 124.5, 116.1 (d, $J_{C\text{–}F}$ = 23.1 Hz), 100.7; HRMS (ESI) *m/z* = 321.0788 calcd for $C_{17}H_{10}FN_4O_2$ [M–H]$^-$, found: 321.0788.

**5-(4-Bromophenyl)-1-phenyl-1*H*-pyrazolo[3,4-*d*]pyrimidine-4,6(5*H*,7*H*)-dione (5c)**

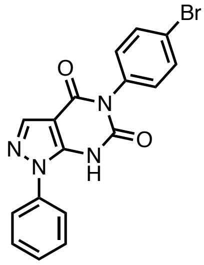

**5c**

1-Bromo-4-isocyanatobenzene was used as the aryl isocyanate. Compound **5c** was obtained as a white solid (39.9 mg, 42% yield). $^{1}$H NMR (600 MHz, DMSO-*d*$_6$) δ 12.50 (brs, 1H), 8.16 (s, 1H), 7.68 (d, *J* = 8.4 Hz, 2H), 7.62 (d, *J* = 7.2 Hz, 2H), 7.58 (t, *J* = 7.2 Hz, 2H), 7.50 (t, *J* = 7.2 Hz, 1H), 7.26 (d, *J* = 8.4 Hz, 2H); $^{13}$C NMR (150 MHz, DMSO-*d*$_6$) δ 158.2, 151.4, 144.0, 138.1, 137.2, 135.9, 132.3, 132.1, 130.0, 129.1, 124.9, 121.6, 100.8; HRMS (ESI) *m/z* = 380.9987 calcd for $C_{17}H_{10}BrN_4O_2$ [M–H]$^-$, found: 380.9994.

**5-(4-Iodophenyl)-1-phenyl-1*H*-pyrazolo[3,4-*d*]pyrimidine-4,6(5*H*,7*H*)-dione (5d)**

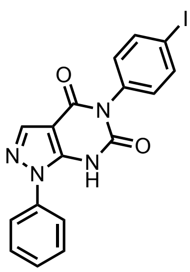

**5d**

1-Iodo-4-isocyanatobenzene was used as the aryl isocyanate. Compound **5d** was obtained as a white solid (35.2 mg, 33% yield). $^{1}$H NMR (600 MHz, DMSO-*d*$_6$) δ 12.49 (brs, 1H), 8.17 (s, 1H), 7.84 (d, *J* = 8.4 Hz, 2H), 7.62–7.55 (m, 4H), 7.51 (t, *J* = 7.2 Hz, 1H), 7.11 (d, *J* = 8.4 Hz,

2H); $^{13}$C NMR (150 MHz, DMSO-$d_6$) δ 158.1, 151.2, 143.5, 138.2, 138.1, 137.0, 136.3, 132.2, 130.0, 129.2, 125.0, 100.8, 94.8; HRMS (ESI) *m/z* = 428.9848 calcd for $C_{17}H_{10}IN_4O_2$ [M–H]$^-$, found: 428.9848.

**1-Phenyl-5-(*p*-tolyl)-1*H*-pyrazolo[3,4-*d*]pyrimidine-4,6(5*H*,7*H*)-dione (5e)**

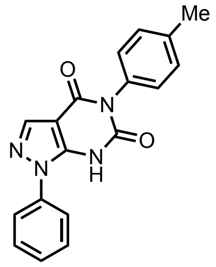


**5e**

1-Isocyanato-4-methylbenzene was used as the aryl isocyanate. Compound **5e** was obtained as a white solid (45.2 mg, 57% yield). $^1$H NMR (600 MHz, DMSO-$d_6$) δ 12.40 (brs, 1H), 8.16 (s, 1H), 7.62 (d, $J$ = 7.8 Hz, 2H), 7.57 (dd, $J$ = 7.8, 7.2 Hz, 2H), 7.51 (t, $J$ = 7.2 Hz, 1H), 7.27 (d, $J$ = 7.8 Hz, 2H), 7.14 (d, $J$ = 7.8 Hz, 2H), 2.37 (s, 3H); $^{13}$C NMR (150 MHz, DMSO-$d_6$) δ 158.3, 151.4, 143.5, 138.1, 137.9, 137.1, 133.7, 130.0, 129.8, 129.4, 129.1, 125.0, 100.9, 21.2; HRMS (ESI) *m/z* = 317.1039 calcd for $C_{18}H_{13}N_4O_2$ [M–H]$^-$, found: 317.1039.

**5-(4-Methoxyphenyl)-1-phenyl-1*H*-pyrazolo[3,4-*d*]pyrimidine-4,6(5*H*,7*H*)-dione (5f)**

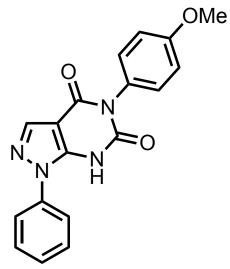


**5f**

1-Isocyanato-4-methoxybenzene was used as the aryl isocyanate. Compound **5f** was obtained as a white solid (6.7 mg, 8% yield). $^1$H NMR (600 MHz, DMSO-$d_6$) δ 12.39 (brs, 1H), 8.14 (s, 1H), 7.62 (d, $J$ = 7.8 Hz, 2H), 7.57 (t, $J$ = 7.8 Hz, 2H), 7.50 (t, $J$ = 7.8 Hz, 1H), 7.17 (d, $J$ = 9.0 Hz, 2H), 7.01 (d, $J$ = 9.0 Hz, 2H), 3.80 (s, 3H); $^{13}$C NMR (150 MHz, DMSO-$d_6$) δ 159.2, 158.5, 151.7, 143.8, 138.1, 137.2, 130.7, 129.9, 129.1, 128.9, 124.9, 114.5, 100.9, 55.8; HRMS (ESI) *m/z* = 333.0988 calcd for $C_{18}H_{13}N_4O_3$ [M–H]$^-$, found: 333.0989.

**1-Phenyl-5-(4-(trifluoromethyl)phenyl)-1*H*-pyrazolo[3,4-*d*]pyrimidine-4,6(5*H*,7*H*)-dione (5g)**

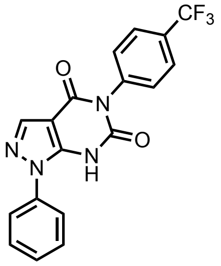

**5g**

1-Isocyanato-4-(trifluoromethyl)benzene was used as the aryl isocyanate. Compound **5g** was obtained as a white solid (24.6 mg, 26% yield). $^{1}$H NMR (600 MHz, DMSO-$d_6$) δ 8.05 (s, 1H), 7.88–7.78 (m, 4H), 7.53 (t, $J$ = 7.8 Hz, 2H), 7.49 (d, $J$ = 7.2 Hz, 2H), 7.40 (t, $J$ = 7.8 Hz, 1H), one proton (NH) was not detected; $^{13}$C NMR (150 MHz, DMSO-$d_6$) δ 159.0, 153.6, 148.9, 141.7, 138.4, 137.6, 131.0, 129.7, 128.5 (q, $J_{C-F}$ = 31.5 Hz), 127.7, 126.2, 124.7 (q, $J_{C-F}$ = 270.0 Hz) 123.3, 100.4; HRMS (ESI) $m/z$ = 371.0756 calcd for $C_{18}H_{10}F_3N_4O_2$ [M–H]$^-$, found: 371.0755.

**5-(3-Fluorophenyl)-1-phenyl-1*H*-pyrazolo[3,4-*d*]pyrimidine-4,6(5*H*,7*H*)-dione (5h)**

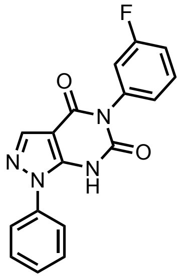

**5h**

1-Fluoro-3-isocyanatobenzene was used as the aryl isocyanate. Compound **5h** was obtained as a white solid (12.4 mg, 15% yield). $^{1}$H NMR (600 MHz, DMSO-$d_6$) δ 8.07 (s, 1H), 7.77 (d, $J$ = 7.8 Hz, 2H), 7.54 (t, $J$ = 7.8 Hz, 2H), 7.49 (m, 1H), 7.43 (t, $J$ = 7.8 Hz, 1H), 7.24 (td, $J$ = 8.4, 2.4 Hz, 1H), 7.17 (m, 1H), 7.10 (d, $J$ = 7.8 Hz, 1H) one proton (NH) was not detected; $^{13}$C NMR (150 MHz, DMSO-$d_6$) δ 162.4 (d, $J_{C-F}$ = 241.2 Hz), 158.6, 152.8, 147.2, 139.0, 137.9, 137.7, 130.4, 129.6, 127.9, 126.2, 123.6, 117.1 (d, $J_{C-F}$ = 23.0 Hz), 114.9 (d, $J_{C-F}$ = 23.1 Hz), 100.4; HRMS (ESI) $m/z$ = 321.0788 calcd for $C_{17}H_{10}FN_4O_2$ [M–H]$^-$, found: 321.0787.

**5-(3-Bromophenyl)-1-phenyl-1*H*-pyrazolo[3,4-*d*]pyrimidine-4,6(5*H*,7*H*)-dione (5i)**

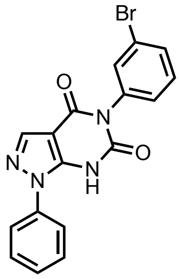

**5i**

1-Bromo-3-isocyanatobenzene was used as the aryl isocyanate. Compound **5i** was obtained as a white solid (26.7 mg, 28% yield). $^{1}$H NMR (600 MHz, DMSO-$d_6$) δ 8.07 (s, 1H), 7.79 (d, $J$ = 7.8 Hz, 2H), 7.59 (d, $J$ = 7.8 Hz, 1H), 7.54 (t, $J$ = 7.8 Hz, 2H), 7.50 (t, $J$ = 1.8 Hz, 1H), 7.45–7.39 (m, 2H), 7.27 (d, $J$ = 7.8 Hz, 1H), one proton (NH) was not detected; $^{13}$C NMR (150 MHz, DMSO-$d_6$) δ 158.8, 153.2, 147.7, 139.1, 138.1, 137.7, 132.8, 131.0, 129.7, 129.3, 128.0, 123.7, 121.4, 100.5, one peak is missing due to overlapping; HRMS (ESI) $m/z$ = 380.9987 calcd for $C_{17}H_{10}BrN_4O_2$ [M–H]$^-$, found: 380.9991.

**5-(2-Fluorophenyl)-1-phenyl-1*H*-pyrazolo[3,4-*d*]pyrimidine-4,6(5*H*,7*H*)-dione (5j)**

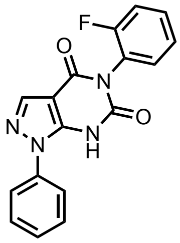

**5j**

1-Fluoro-2-isocyanatobenzene was used as the aryl isocyanate. Compound **5j** was obtained as a white solid (32.7 mg, 41% yield). $^{1}$H NMR (600 MHz, DMSO-$d_6$) δ 8.18 (s, 1H), 7.66 (d, $J$ = 7.2 Hz, 2H), 7.58 (t, $J$ = 7.2 Hz, 2H), 7.52–7.47 (m, 2H), 7.43 (td, $J$ = 7.8, 1.8 Hz, 1H), 7.38 (m, 1H), 7.32 (td, $J$ = 7.8, 1.8 Hz, 1H), one proton (NH) was not detected; $^{13}$C NMR (150 MHz, DMSO-$d_6$) δ 158.4 (d, $J_{C–F}$ = 245.7 Hz), 157.6, 151.0, 144.4, 138.1, 137.2, 132.1, 130.9 (d, $J_{C–F}$ = 7.1 Hz), 129.9, 129.1, 125.2, 125.0, 123.8 (d, $J_{C–F}$ = 14.4 Hz), 116.4 (d, $J_{C–F}$ = 18.8 Hz), 100.4; HRMS (ESI) $m/z$ = 321.0788 calcd for $C_{17}H_{10}FN_4O_2$ [M–H]$^-$, found: 321.0787.

**5-(2-Chlorophenyl)-1-phenyl-1*H*-pyrazolo[3,4-*d*]pyrimidine-4,6(5*H*,7*H*)-dione (5k)**

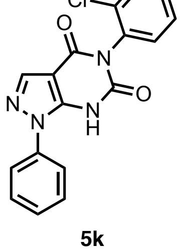

**5k**

1-Chloro-2-isocyanatobenzene was used as the aryl isocyanate. Compound **5k** was obtained as a white solid (25.5 mg, 30% yield). $^{1}$H NMR (600 MHz, DMSO-$d_6$) δ 8.20 (s, 1H), 7.66–7.62 (m, 3H), 7.58 (t, $J$ = 7.2 Hz, 2H), 7.51 (t, $J$ = 7.2 Hz, 1H), 7.50–7.46 (m, 3H), one proton (NH) was not detected; $^{13}$C NMR (150 MHz, DMSO-$d_6$) δ 157.5, 150.7, 144.0, 138.2, 137.0, 134.1, 132.8, 132.0, 130.7, 130.1, 130.0, 129.3, 128.5, 125.2, 100.5; HRMS (ESI) $m/z$ = 337.0492 calcd for $C_{17}H_{10}ClN_4O_2$ $[M–H]^-$, found: 337.0492.

**5-(2-Bromophenyl)-1-phenyl-1*H*-pyrazolo[3,4-*d*]pyrimidine-4,6(5*H*,7*H*)-dione (5l)**

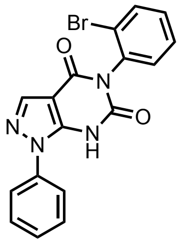


**5l**

1-Bromo-2-isocyanatobenzene was used as the aryl isocyanate. Compound **5l** was obtained as a white solid (11.8 mg, 12% yield). $^{1}$H NMR (600 MHz, DMSO-$d_6$) δ 8.21 (s, 1H), 7.78 (d, $J$ = 7.8 Hz, 1H), 7.64 (d, $J$ = 7.8 Hz, 2H), 7.58 (t, $J$ = 7.8 Hz, 2H), 7.52 (t, $J$ = 7.8 Hz, 2H), 7.47 (d, $J$ = 7.8 Hz, 1H), 7.40 (td, $J$ = 7.8, 1.8 Hz, 1H), one proton (NH) was not detected; $^{13}$C NMR (150 MHz, DMSO-$d_6$) δ 157.4, 150.6, 143.9, 138.2, 137.0, 135.7, 133.2, 132.0, 130.9, 130.0, 129.3, 129.1, 125.2, 123.6, 100.6; HRMS (ESI) $m/z$ = 380.9987 calcd for $C_{17}H_{10}BrN_4O_2$ $[M–H]^-$, found: 380.9994.

**5-(3,4-Difluorophenyl)-1-phenyl-1*H*-pyrazolo[3,4-*d*]pyrimidine-4,6(5*H*,7*H*)-dione (5m)**

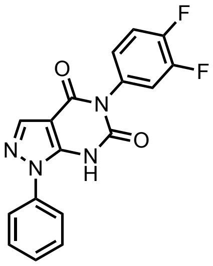


**5m**

1,2-Difluoro-4-isocyanatobenzene was used as the aryl isocyanate. Compound **5m** was obtained as a white solid (34.8 mg, 41% yield). $^{1}$H NMR (600 MHz, DMSO-$d_6$) δ 8.13 (s, 1H), 7.69 (d, $J$ = 7.2 Hz, 2H), 7.59–7.45 (m, 5H), 7.18 (d, $J$ = 9.0 Hz, 1H), one proton (NH) was not detected; $^{13}$C NMR (150 MHz, DMSO-$d_6$) δ 158.4, 152.2, 149.7 (dd, $J_{C–F}$ = 244.2, 12.9 Hz), 149.6 (dd, $J_{C–F}$ = 244.1, 13.1 Hz), 145.6, 138.0, 137.5, 133.6 (d, $J_{C–F}$ = 5.7 Hz), 129.9, 128.7, 127.2 (d, $J_{C–F}$ = 2.9 Hz), 124.4, 119.5 (d, $J_{C–F}$ = 18.6 Hz), 117.8 (d, $J_{C–F}$ = 17.3 Hz), 100.6; HRMS (ESI) $m/z$ = 339.0694 calcd for $C_{17}H_9F_2N_4O_2$ $[M–H]^-$, found: 339.0690.

**5-(3-Chloro-4-fluorophenyl)-1-phenyl-1*H*-pyrazolo[3,4-*d*]pyrimidine-4,6(5*H*,7*H*)-dione (5n)**

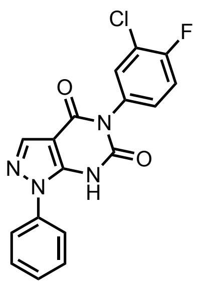

**5n**

2-Chloro-1-fluoro-4-isocyanatobenzene was used as the aryl isocyanate. Compound **5n** was obtained as a white solid (37.2 mg, 42% yield). $^{1}$H NMR (600 MHz, DMSO-$d_6$) δ 12.55 (brs, 1H), 8.17 (s, 1H), 7.64 (dd, $J$ = 7.2, 1.8 Hz, 1H), 7.63–7.56 (m, 4H), 7.55–7.49 (m, 2H), 7.36 (m, 1H); $^{13}$C NMR (150 MHz, DMSO-$d_6$) δ 158.2, 157.3 (d, $J_{C–F}$ = 245.9 Hz), 151.4, 144.0, 138.1, 137.1, 133.5, 132.1, 130.9 (d, $J_{C–F}$ = 7.2 Hz), 130.0, 129.2, 124.9, 119.9 (d, $J_{C–F}$ = 18.8 Hz), 117.5 (d, $J_{C–F}$ = 23.0 Hz), 100.8; HRMS (ESI) $m/z$ = 355.0398 calcd for $C_{17}H_9ClFN_4O_2$ $[M–H]^-$, found: 355.0398.

**5-(3-Chloro-4-methylphenyl)-1-phenyl-1*H*-pyrazolo[3,4-*d*]pyrimidine-4,6(5*H*,7*H*)-dione (5o)**

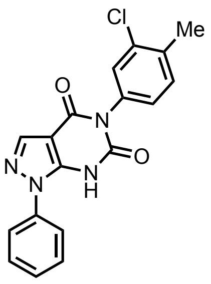

**5o**

2-Chloro-4-isocyanato-1-methylbenzene was used as the aryl isocyanate. Compound **5o** was obtained as a white solid (12.7 mg, 14% yield). $^{1}$H NMR (600 MHz, DMSO-$d_6$) δ 12.48 (brs, 1H), 8.16 (s, 1H), 7.62 (d, $J$ = 7.8 Hz, 2H), 7.58 (t, $J$ = 7.8 Hz, 2H), 7.50 (t, $J$ = 7.8 Hz, 1H), 7.45 (d, $J$ = 7.8 Hz, 1H), 7.40 (s, 1H), 7.17 (d, $J$ = 7.8 Hz, 1H) 2.38 (s, 3H); $^{13}$C NMR (150 MHz, DMSO-$d_6$) δ 158.2, 151.5, 144.0, 138.1, 137.2, 135.9, 135.4, 133.4, 131.7, 130.1, 130.0, 129.1, 128.6, 124.9, 100.8, 19.8; HRMS (ESI) $m/z$ = 351.0649 calcd for $C_{18}H_{12}ClN_4O_2$ $[M–H]^-$, found: 351.0649.

**5-(2,4-Difluorophenyl)-1-phenyl-1*H*-pyrazolo[3,4-*d*]pyrimidine-4,6(5*H*,7*H*)-dione (5p)**

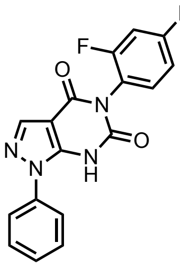

**5p**

2,4-Difluoro-1-isocyanatobenzene was used as the aryl isocyanate. Compound **5p** was obtained as a white solid (51.2 mg, 60% yield). $^{1}$H NMR (600 MHz, DMSO-$d_6$) δ 8.13 (s, 1H), 7.76 (d, $J$ = 7.8 Hz, 2H), 7.55 (t, $J$ = 7.8 Hz, 2H), 7.51–7.38 (m, 3H), 7.20 (t, $J$ = 8.4 Hz, 1H), one proton (NH) was not detected; $^{13}$C NMR (150 MHz, DMSO-$d_6$) δ 162.3 (dd, $J_{C–F}$ = 245.6, 11.6 Hz), 158.6 (dd, $J_{C–F}$ = 249.9, 13.1 Hz), 158.1, 152.1, 146.8, 137.9, 137.8, 133.2 (d, $J_{C–F}$ = 10.1 Hz), 129.8, 128.4, 124.3, 121.1 (d, $J_{C–F}$ = 10.1 Hz), 112.2 (d, $J_{C–F}$ = 23.0 Hz), 105.0 (t, $J_{C–F}$ = 24.5 Hz), 100.2; HRMS (ESI) $m/z$ = 339.0694 calcd for $C_{17}H_9F_2N_4O_2$ $[M–H]^-$, found:339.0692.

**5-(4-Bromo-2-fluorophenyl)-1-phenyl-1*H*-pyrazolo[3,4-*d*]pyrimidine-4,6(5*H*,7*H*)-dione (5q)**

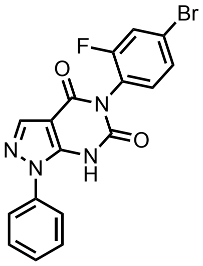

**5q**

4-Bromo-2-fluoro-1-isocyanatobenzene was used as the aryl isocyanate. Compound **5q** was obtained as a white solid (42.1 mg, 42% yield). $^{1}$H NMR (600 MHz, DMSO-$d_6$) δ 8.20 (s, 1H), 7.77 (dd, $J$ = 9.0, 1.8 Hz, 1H), 7.63 (m, 2H), 7.61–7.54 (m, 3H), 7.51 (t, $J$ = 7.2 Hz, 1H), 7.44 (t, $J$ = 8.4 Hz, 1H), one proton (NH) was not detected; $^{13}$C NMR (150 MHz, DMSO-$d_6$) δ 158.4 (d, $J_{C–F}$ = 251.4 Hz), 157.4, 150.6, 144.0, 138.1, 137.0, 133.7, 130.0, 129.3, 128.6, 125.2, 123.4 (d, $J_{C–F}$ = 12.9 Hz), 122.5 (d, $J_{C–F}$ = 10.1 Hz), 120.0 (d, $J_{C–F}$ = 23.1 Hz), 100.4; HRMS (ESI) $m/z$ = 398.9893 calcd for $C_{17}H_9BrFN_4O_2$ $[M–H]^-$, found: 398.9893.

**5-(2,5-Dichlorophenyl)-1-phenyl-1*H*-pyrazolo[3,4-*d*]pyrimidine-4,6(5*H*,7*H*)-dione (5r)**

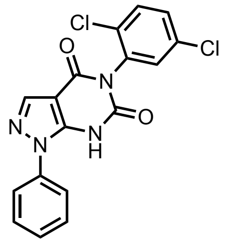

**5r**

1,4-Dichloro-2-isocyanatobenzene was used as the aryl isocyanate. Compound **5r** was obtained as a white solid (68.5 mg, 73% yield). $^{1}$H NMR (600 MHz, DMSO-$d_6$) δ 8.12 (s, 1H), 7.79 (d, *J* = 7.2 Hz, 2H), 7.68–7.61 (m, 2H), 7.58–7.51 (m, 3H), 7.44 (t, *J* = 7.2 Hz, 1H), one proton (NH) was not detected; $^{13}$C NMR (150 MHz, DMSO-$d_6$) δ 158.0, 152.2, 147.6, 137.9, 137.8, 136.5, 132.2, 132.1, 132.0, 131.3, 130.3, 129.8, 128.3, 124.0, 100.2; HRMS (ESI) *m/z* = 371.0103 calcd for $C_{17}H_9Cl_2N_4O_2$ [M–H]$^-$, found: 371.0102.

**5-(2,3-Dichlorophenyl)-1-phenyl-1*H*-pyrazolo[3,4-*d*]pyrimidine-4,6(5*H*,7*H*)-dione (5s)**

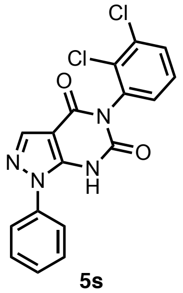

**5s**

1,2-Dichloro-3-isocyanatobenzene was used as the aryl isocyanate. Compound **5s** was obtained as a white solid (23.1 mg, 25% yield). $^{1}$H NMR (600 MHz, DMSO-$d_6$) δ 8.13 (s, 1H), 7.77 (d, *J* = 7.8 Hz, 2H), 7.73 (dd, *J* = 7.8, 1.8 Hz, 1H), 7.55 (t, *J* = 7.8 Hz, 2H), 7.49 (t, *J* = 7.8 Hz, 1H), 7.47–7.43 (m, 2H), one proton (NH) was not detected; $^{13}$C NMR (150 MHz, DMSO-$d_6$) δ 157.9, 151.9, 147.1, 137.9, 137.8, 137.0, 132.4, 131.7, 130.9, 130.8, 129.8, 128.9, 128.4, 124.1, 100.2; HRMS (ESI) *m/z* = 371.0103 calcd for $C_{17}H_9Cl_2N_4O_2$ [M–H]$^-$, found: 371.0102.

**5-(2,4-Dichlorophenyl)-1-phenyl-1*H*-pyrazolo[3,4-*d*]pyrimidine-4,6(5*H*,7*H*)-dione (5t: TU-923)**

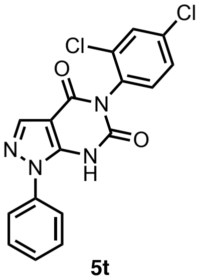

**5t**

2,4-Dichloro-1-isocyanatobenzene was used as the aryl isocyanate. Compound **5t** was obtained as a white solid (53.1 mg, 57% yield). $^1$H NMR (600 MHz, DMSO-$d_6$) δ 8.22 (s, 1H), 7.84 (d, $J$ = 2.4 Hz, 1H), 7.62 (d, $J$ = 7.8 Hz, 2H), 7.61–7.56 (m, 3H), 7.55–7.50 (m, 2H), one proton (NH) was not detected; $^{13}$C NMR (150 MHz, DMSO-$d_6$) δ 157.3, 150.4, 143.8, 138.2, 136.9, 134.5, 134.1, 133.3, 133.2, 130.0, 129.8, 129.4, 128.8, 125.3, 100.5; HRMS (ESI) $m/z$ = 371.0103 calcd for $C_{17}H_9Cl_2N_4O_2$ [M–H]$^-$, found: 371.0104.

**5-(2,6-Dichlorophenyl)-1-phenyl-1*H*-pyrazolo[3,4-*d*]pyrimidine-4,6(5*H*,7*H*)-dione (5u)**

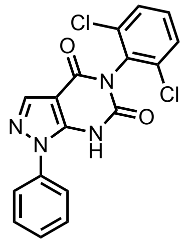

**5u**

1,3-Dichloro-2-isocyanatobenzene was used as the aryl isocyanate. Compound **5u** was obtained as a white solid (45.1 mg, 48% yield). $^1$H NMR (600 MHz, DMSO-$d_6$) δ 8.18 (s, 1H), 7.78 (d, $J$ = 7.2 Hz, 2H), 7.64 (d, $J$ = 7.8 Hz, 2H), 7.55 (t, $J$ = 7.2 Hz, 2H), 7.50 (t, $J$ = 7.8 Hz, 1H), 7.46 (t, $J$ = 7.2 Hz, 1H), one proton (NH) was not detected; $^{13}$C NMR (150 MHz, DMSO-$d_6$) δ 157.1, 151.0, 146.8, 138.0, 137.6, 134.8, 132.8, 131.4, 129.8, 129.1, 128.6, 124.5, 99.9; HRMS (ESI) $m/z$ = 371.0103 calcd for $C_{17}H_9Cl_2N_4O_2$ [M–H]$^-$, found: 371.0100.

**1-Phenyl-5-(2,4,6-trichlorophenyl)-1*H*-pyrazolo[3,4-*d*]pyrimidine-4,6(5*H*,7*H*)-dione (5v)**

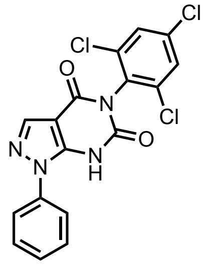

**5v**

1,3,5-Trichloro-2-isocyanatobenzene was used as the aryl isocyanate. Compound **5v** was obtained as a white solid (91.6 mg, 90% yield). $^1$H NMR (600 MHz, DMSO-$d_6$) δ 8.26 (s, 1H), 7.92 (s, 2H), 7.66 (d, $J$ = 7.8 Hz, 2H), 7.58 (t, $J$ = 7.8 Hz, 2H), 7.52 (t, $J$ = 7.8 Hz, 1H), one proton (NH) was not detected; $^{13}$C NMR (150 MHz, DMSO-$d_6$) δ 156.4, 149.6, 144.1, 138.3, 136.9, 135.7, 135.3, 131.3, 130.0, 129.5, 129.2, 125.5, 100.0; HRMS (ESI) $m/z$ = 404.9713 calcd for $C_{17}H_8Cl_3N_4O_2$ [M–H]$^-$, found: 404.9716.

**[1]H NMR (600 MHz, DMSO-$d_6$) of TU-892:**

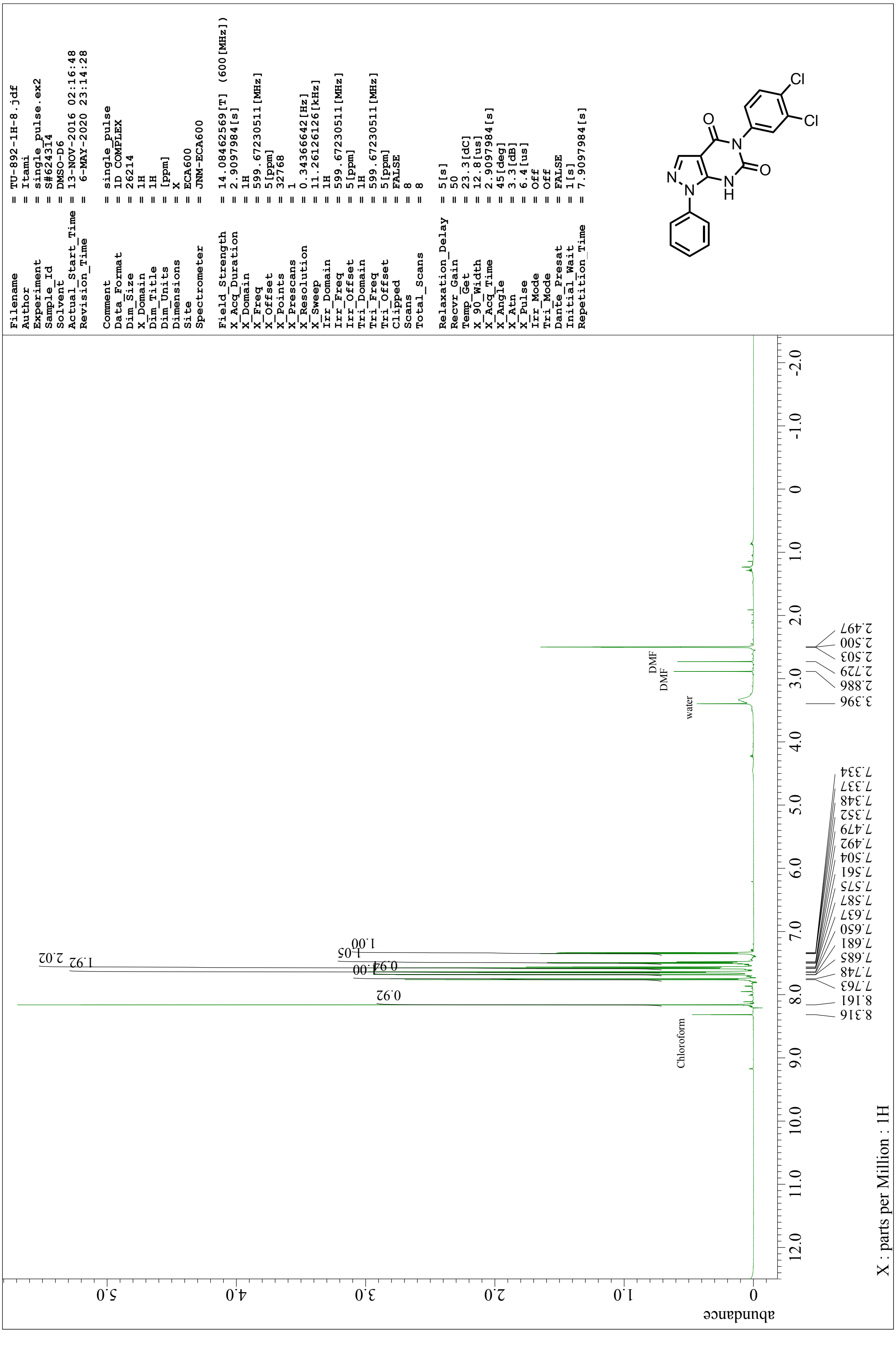

**$^{13}$C NMR (150 MHz, DMSO-$d_6$) of TU-892:**

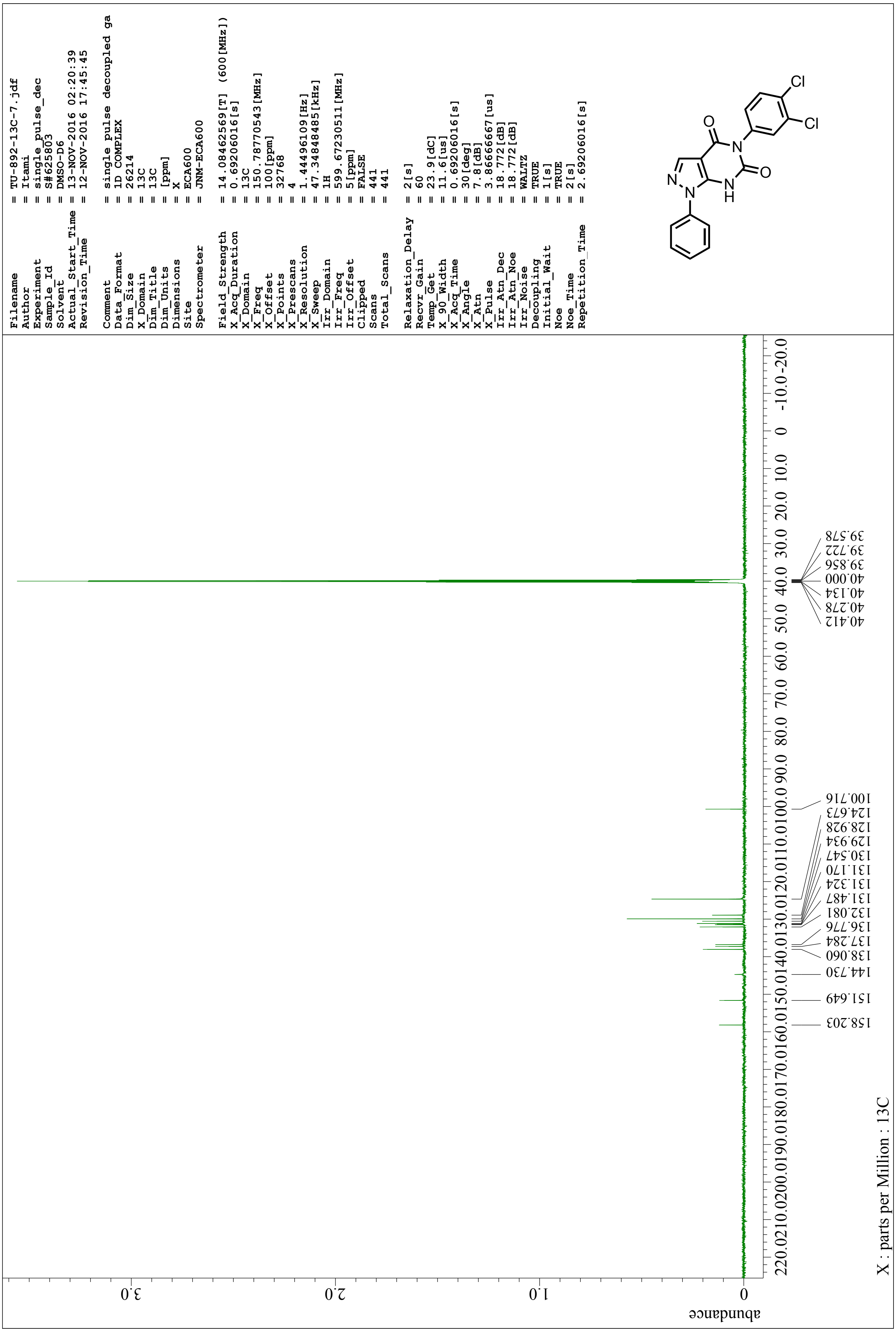

**[1]H NMR (600 MHz, DMSO-$d_6$) of TU-923:**

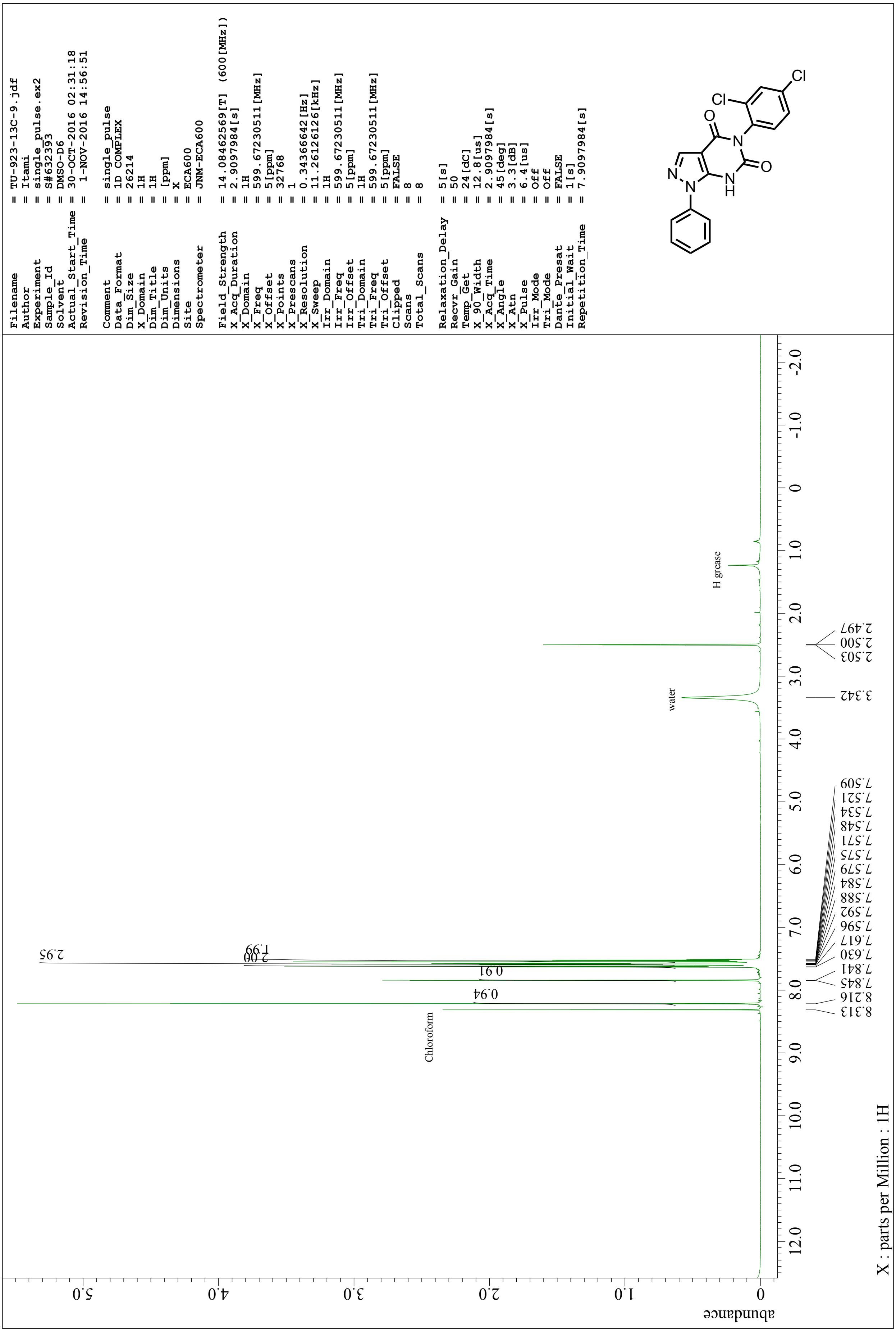

[13]C NMR (150 MHz, DMSO-$d_6$) of TU-923:

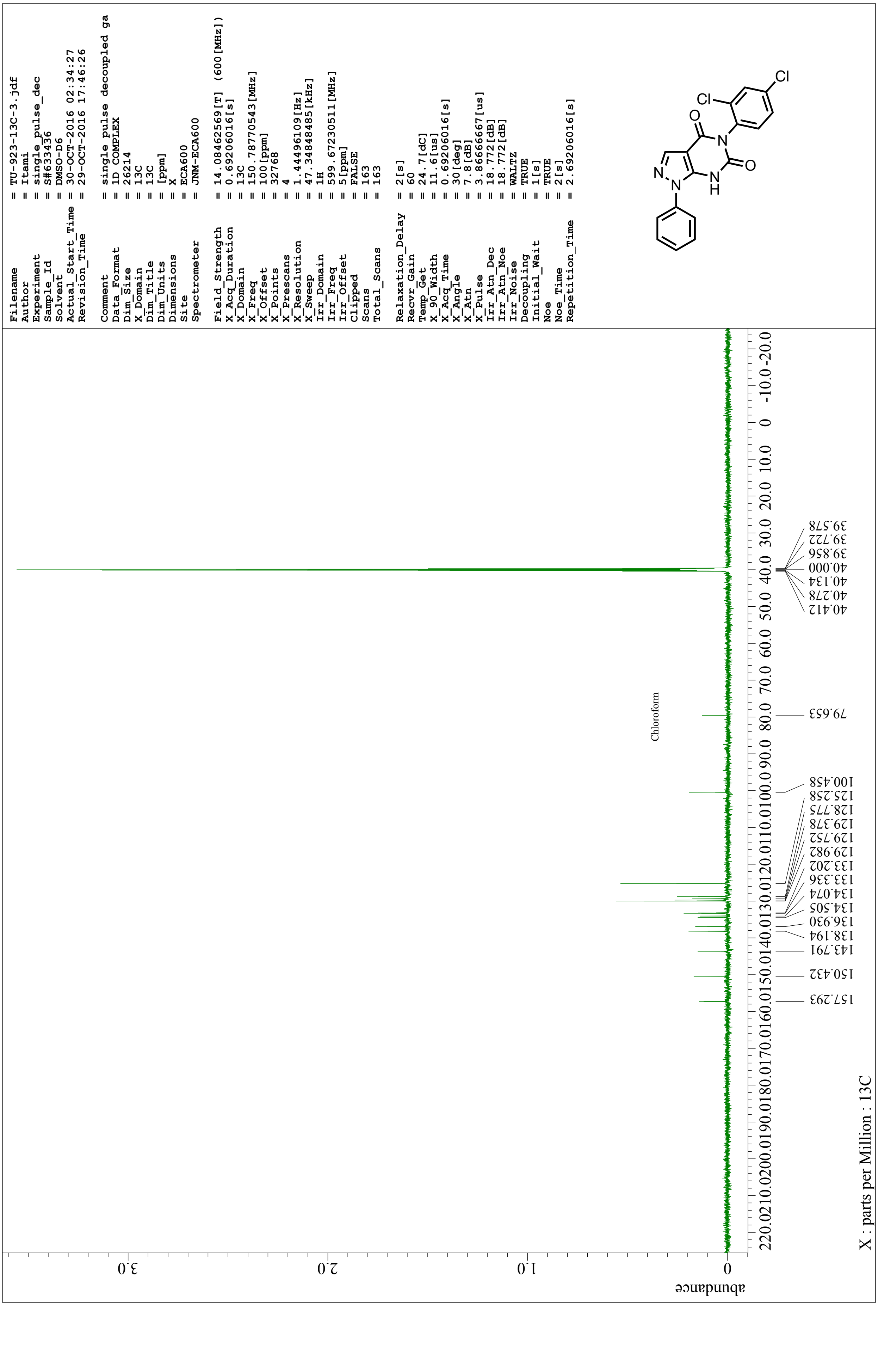
